# SeedMatExplorer: The transcriptome atlas of Arabidopsis seed maturation

**DOI:** 10.1101/2024.11.04.621888

**Authors:** Mariana A. S. Artur, Robert A. Koetsier, Leo A. J. Willems, Lars L. Bakermans, Annabel D. van Driel, Joram A. Dongus, Bas J. W. Dekkers, Alexandre C. S. S. Marques, Asif Ahmed Sami, Harm Nijveen, Leónie Bentsink, Henk Hilhorst, Renake Nogueira Teixeira

**Affiliations:** Laboratory of Plant Physiology, Wageningen Seed Science Centre, Wageningen University & Research, Wageningen, 6708 PB, The Netherlands; Bioinformatics Group, Wageningen University & Research, Wageningen, 6708 PB, The Netherlands; Department of Molecular and Cell Biology, University of Cape Town, Cape Town, South Africa; Hazera Seeds B.V., Schanseind 27, 4921 PM Made

**Keywords:** seed maturation, gene regulatory networks, transcriptomics, desiccation tolerance, longevity

## Abstract

**Background:** Seed maturation is a critical developmental phase during which seeds acquire traits essential for nutritional value, desiccation tolerance, and long-term survival. Abscisic acid (ABA) signalling is a key regulator of this process, coordinating gene expression programs underlying the acquisition of seed quality traits. However, the molecular regulation of many of these traits remains poorly understood. To address this, we performed a comprehensive analysis of seed maturation in *Arabidopsis thaliana*, combining physiological and transcriptomic approaches across wild-type plants and mutants affected in ABA biosynthesis, signalling, and catabolism.

**Results:** We generated a high-resolution transcriptome dataset covering seed development from 12 days after pollination to the dry seed stage in wild-type and ten mutant lines. In parallel, we characterized the temporal acquisition of multiple seed traits, including germination capacity, dormancy, chlorophyll fluorescence, longevity and desiccation tolerance. Integration of these datasets using weighted gene co-expression network analysis (WGCNA) identified gene modules associated with specific trait acquisition patterns. This approach enabled the identification of coordinated transcriptional programs linked to distinct seed quality traits, extending beyond individual gene-level analyses. Notably, modules associated with desiccation tolerance and longevity were enriched for genes involved in stress responses and ABA-regulated pathways, highlighting the complex and multifactorial regulation of these traits.

**Conclusions:** This study provides a comprehensive physiological and transcriptomic framework for understanding seed maturation and the acquisition of key seed quality traits in *Arabidopsis thaliana*. By linking gene expression dynamics to trait development, our work offers new insights into the regulatory networks underlying seed resilience and storage capacity. The dataset is made accessible through SeedMatExplorer (https://www.bioinformatics.nl/SeedMatExplorer), an open-access web platform that enables interactive exploration and supports hypothesis generation. Together, this resource represents a valuable tool for advancing research on seed biology and improving seed performance in agricultural contexts.

## Background

Seeds are a crucial stage of a plant’s life cycle, allowing the plant species to survive in space and time, and through severe environmental challenges. In most plant species, seed development can be divided into embryogenesis, maturation, and drying. During embryogenesis (also called morphogenesis or histodifferentiation) cell division leads to the embryo body plan (1, 2). In the maturation phase (also called seed filling), cell division ceases, cells expand due to nutrient reserve deposition, and embryonic chlorophyll is gradually degraded (3, 4). In most species, a drying phase (or late seed maturation) takes place after seed filling, which is marked by a decrease in the seed moisture content leading to a quiescent state (4–8). Seed maturation has gained increasing attention as a phase in which key seed quality traits (SQTs) are acquired, and has been proposed as an evolutionary landscape for the emergence of novel genes promoting functional specialization contributing to survival, resilience, and storage (9). Seed longevity, the capacity of seeds to remain alive during storage, is a critical SQT for *ex situ* conservation of plant genetic resources and for seed persistence in agricultural and ecological contexts (4, 10, 11). Dormancy and desiccation tolerance (DT) are two other SQTs of large agronomical and ecological importance associated with successful seedling establishment (12) and plant adaptation on land (13), respectively.

During maturation, the acquisition of the above-mentioned SQTs requires the coordinated activity of multiple genes. The hormone abscisic acid (ABA) is a crucial positive regulator of several of these genes (14–21), and accumulates during maturation and declines during drying (22–24). With the isolation of mutants deficient in and insensitive to ABA in the early ‘80s and ‘90s, *Arabidopsis thaliana* became one of the main models to study the genetic and molecular control of SQTs during maturation (22, 25–29). ABA deficiency leads to absence of primary dormancy, and plants with overexpression or loss of ABA biosynthesis genes have increased and decreased seed dormancy, respectively (22, 27, 29–31). Mutations of ABA catabolism genes, such as the enzyme *CYP707A2*, lead to ABA over-accumulation and enhance dormancy (32, 33). Furthermore, mutants with strong defects in ABA sensitivity fail to acquire DT (34, 35). *ABSCISIC ACID INSENSITIVE 3* (*ABI3*) is one of the main master regulators of seed maturation and of multiple SQTs (22, 25, 26, 36–38), being one of the main transducers of the ABA signal during seed maturation. Strong mutant alleles of *ABI3* (e.g., *abi3-4*; *abi3-5* and *abi3-6*) fail to degrade chlorophyll and to acquire dormancy, DT and longevity, and also fail to accumulate seed storage proteins (SSPs), Raffinose Family Oligosaccharides (RFOs) and late embryogenesis abundant proteins (LEAs) (34–36, 39, 40).

Advances in gene expression analyses, particularly transcriptomics, have facilitated genetic studies further elucidating ABI3-mediated transcriptional control of seed maturation. Using a strong *ABI3* allele mutant (*abi3-6*), Delmas, Sankaranarayanan and colleagues (41) showed that ABI3 regulates embryo degreening by controlling chlorophyll degradation enzymes. Studies on the weak *ABI3* allele (*abi3-1*) identified the green-seed (GRS) locus and *DELAY OF GERMINATION* (*DOG*) 1 as enhancers of the *ABI3*-dependent green seed phenotype (42, 43). *DOG1* is a key regulator of seed dormancy in Arabidopsis (44–46) and appears to act largely independently of ABA, although their pathways converge on downstream maturation processes (43, 46, 47). *DOG1* mutants show reduced longevity and lower expression of maturation genes such as *LEAs* and *SSPs*, while *abi3-1 dog1-1* double mutants produce ABA-insensitive, green seeds (43, 45), highlighting the roles of *DOG1* and *ABI3* in seed maturation and chlorophyll degradation (41–43, 48–50). Integration of transcriptomics with molecular assays such as chromatin immunoprecipitation (ChIP) has further identified direct and indirect ABI3 targets involved in seed maturation, desiccation tolerance, dormancy, and longevity (51, 52). Together, these studies highlight the complex regulation of SQT acquisition during seed maturation, although how these traits are coordinated and interconnected remains poorly understood.

Gene co-expression network analysis has been a useful method to study the regulation of complex traits, and has been widely used to study the regulation of seed maturation across species (53–69). Despite extensive transcriptomics studies of Arabidopsis seed development and advancements on co-expression network analyses (17, 70–75), the regulation of STQ acquisition during maturation, especially the distinct regulatory modules involved in their control, still remains poorly resolved, in part due to overlapping developmental processes that hinder the identification of trait-specific regulatory modules.

To address these challenges, we developed an integrative framework to identify regulators of SQTs in Arabidopsis. Combining detailed physiological analyses of wild-type (Columbia-0) and multiple seed maturation and ABA-related mutants with high-resolution, time-resolved RNA-seq, we constructed gene co-expression networks and performed module-trait correlations to identify the regulatory modules and regulators of specific SQTs. We provide the resulting physiological and transcriptomic resource as an open-access platform, SeedMatExplorer (https://www.bioinformatics.nl/SeedMatExplorer), for further exploitation of our broad datasets by the scientific community.

## Materials and Methods

### Plant materials and growth conditions

*Arabidopsis thaliana* (L.) Heynh. accession Columbia (Col-0) and several mutants known to display maturation defects (Supplemental Table 1) were used. *cyp707a2-1* (Salk_072410, (33)), *aba2-1* (27), *abi3-6* (76), *dog1-4* (SM_3_20808,(77)), *abi3-12* (78) were obtained in the Nottingham Arabidopsis Stock Centre (NASC). The *dog1-4* is an induced transposon insertion mutant (SM_3_20808, (77)) and was obtained as described by (45). Genotyping of T-DNA mutants was performed by standard PCR using primers provided by the Salk Institute Genomic Analysis Laboratory (http://signal.salk.edu/tdnaprimers.2.html). The *abi3-12* described by (78) was genotyped using (d)CAPS markers forward: ATGGCGAGACAGAGGAGGTTCTTGTC and Reverse: CCGAGGTTACCCACGTCGCTTTGCT developed using dCAPS Finder 2.0 (http://helix.wustl.edu/dcaps/dcaps.html) (79) and wild-type fragments were digested with Ava II restriction enzyme (NEB, R0152). Double and triple mutants were obtained after crossing their respective parents. F2 and F3 individuals were genotyped using PCR and CAPS markers to confirm their double or triple mutant genotype. To generate dormant Col-0 lines (Col-0+*DOG1*-Cvi), Col-0 plants were transformed with a 5 kb construct containing the *DOG1*-Cvi gene (45). Tail-PCR located the insertion site in line 2-1-2, RT-qPCR on DNA confirmed the single insertion in this line. Plants were grown on 4 x 4 cm Rockwool blocks in a growth chamber at 20°C/18 °C (day/night) under a 16-h photoperiod of artificial light (150 μmol m^-2^ s^-1^) and 70% relative humidity (RH). Plants were watered three times per week with the standard nutrient solution Hyponex (He et al., 2014). Several flowers were tagged in every plant of all genotypes just before anthesis to allow follow up on days after pollination (DAP). For each round of experiments plants were grown together and seeds were harvested from 12 DAP to 26 DAP in 2 days intervals. For each sample, four biological replicates consisting of three plants were used, and 4 to 6 siliques were collected from these biological replicates at the right DAP. Fresh siliques were immediately dissected, and the seeds were carefully removed for seed quality traits (SQTs) phenotyping. The samples for RNA extraction were immediately frozen in liquid nitrogen and stored at -80 °C. The experiment was repeated 3 times, hereby called rounds 1, 2, and 3. Round 1 consisted of pre-tests to assess scalability for phenotyping and RNA-seq sampling and rounds 2 and 3 consisted of phenotyping for moisture content, fresh weight, dry weight, germination, dormancy, desiccation tolerance (DT), chlorophyll fluorescence, and longevity. RNA-seq sampling was also done for Col-0 and the mutants used in this study in both rounds 2 and 3 (Supplemental Table 2).

### Seed quality trait phenotyping

All seed trait phenotyping analyses were performed using three to four biological replicates, each consisting of three plants as a replicate, from which 4-6 siliques were collected. The water content of the seed samples was determined gravimetrically on a fresh weight basis. Siliques from four biological replicates were collected from any of the plants within the replicate, depending on availability at the appropriate DAP. For each genotype and time point, three siliques were sampled per biological replicate. Seeds were excised from the siliques using fine tweezers, the fresh weight was determined within 10 minutes after excision and the dry weight was determined after 24 hours of drying at 105 °C. To assess germination capacity and dormancy, four replicates of seeds excised from 4-6 siliques were sown on two layers of blue germination paper soaked with 40 mL HLO and incubated at 22 °C under continuous light for 7 days. Images were taken two to three times per day to determine maximum germination percentage (gMax%) and time to 50% germination (T50) using the Germinator program (80). After 7 days, germinated seeds were removed, and the remaining seeds were incubated for an additional 5 days on paper supplemented with 10 μM GALLL and 10 mM KNOL to break dormancy. Germination was calculated as the percentage of seeds germinated after 7 days, and dormancy as the percentage of seeds germinating after the dormancy-breaking treatment. To measure the chlorophyll content we used a non-destructive method based on chlorophyll fluorescence. The seeds were analysed with the PathoScreen system developed by Wageningen University and Research and commercialised by PhenoVation

B.V. (81). The same four replicates of seeds from 4-6 siliques used for germination phenotyping were analysed in the PathoScreen and the chlorophyll fluorescence signals were recorded as the ratio between the variable fluorescence (fv) and the maximum fluorescence. DT was assessed for Col-0 according to Ooms et al. (1993). In summary, four replicates of seeds from 4-6 siliques at different DAPs were dissected and artificially desiccated for 48 hours at 22 °C and 30% relative humidity (RH) in the dark. Desiccated seeds were sown onto two layers of blue paper soaked with 40 ml of demi water and DT was calculated as the percentage of normal seedling formation after 7 days of germination. Longevity assessment was performed using a controlled deterioration test (CDT) for Col-0 from 16 – 26 DAP (82, 83) and Elevated Partial Pressure of Oxygen (EPPO) assay for mature seeds of all genotypes (84). For CDT, four replicates of Col-0 seeds from 3-4 siliques were placed in an opened 1.5ml tube and stored above a saturated KCl solution in a closed ventilated tank at 80−85 % relative humidity (RH) and 40 °C for 2 and 5 days. RH was monitored using a data logger (EL-USB-1-LCB, Lascar Electronics). For the EPPO assay three replicates of seeds from the total bulk harvest (freshly harvested mature dry seeds) from all genotypes were placed in 2 ml tubes perforated with a hole of approximately 1 mm diameter closed with an oxygen-permeable polyethylene membrane, placed in 1.5 L steel tanks, filled with air at a rate of 0.4 MPa per minute from buffer tanks until the tank pressure reached approximately 20 MPa (85). The tanks were stored at 22°C and 55% RH, and the seeds were collected after 21, 42, 62, 77, 91, 105 and 119 days. After CDT and EPPO, seed germination data analysis was performed as described previously using Germinator (80). ABA quantification was performed on 18 DAP and dry seeds of all genotypes. In summary, five milligrams of seeds from four biological replicates were ground to fine powder and extracted with 1 mL of 10% (v/v) methanol containing 100 nM deuterated ABA as an internal standard. The extraction was further carried out according to (86) with minor modifications, namely the use of a Strata-X 30 mg/3 mL SPE column (phenomenex). For the detection and quantification of ABA by LC-MS/MS, we used the same procedure as (87) with minor modifications. Multiple reaction monitoring (MRM) mode was used for the quantification and identification of ABA, by comparing the retention time and MRM transitions (263.25>153.15 + 263.25>219.15). The final content of each sample was normalised using the internal standard.

### RNA extraction, sequencing, and analysis

Total RNA was extracted using a modified hot borate protocol (88). Two to three biological replicates were used per treatment (genotype and time point), each consisting of three plants.

For each replicate, seeds were collected from four siliques sampled from these plants, depending on availability at the corresponding developmental stage. The siliques within a biological replicate were pooled prior to RNA extraction, resulting in a total of 123 samples (Supplemental Table 2). The samples were homogenised and mixed with 800 μL of extraction buffer heated to 80 °C (0.2 M Na borate decahydrate (Borax), 30 mM EGTA, 1% SDS, 1% Na deoxy-cholate (Na-DOC), 1.6 mg DTT and 48 mg PVP40). One milligram of proteinase K was added to this suspension and incubated for 15 min at 42°C. After adding 64 μL of 2 M KCl, the samples were incubated on ice for 30 min and subsequently centrifuged for 20 min at 12,000 g. 270 μL of ice-cold 8 M LiCl was added to the supernatant in a final concentration of 2 M and the tubes were incubated overnight on ice. After centrifugation for 20 min at 12,000 g at 4 °C, the pellets were washed with 750 μL ice-cold 2 M LiCl. The samples were centrifuged for 10 min at 10,000 g at 4 °C and the pellets were resuspended in 100 μL DEPC treated water. The samples were cleaned with phenol chloroform and treated with DNAse (RQ1 DNase, Promega). The RNA quality and concentration were assessed by agarose gel electrophoresis and UV spectrophotometry. RNA was processed for use in RNA-seq with mRNA enrichment polyA capture (Illumina Incorporated, San Diego, CA, USA). The sequencing of a total of 123 samples was performed strand specific on Illumina Hiseq 2500, using cDNA and random hexamer priming, and generating single-end 125 nt reads. The raw reads were mapped to the TAIR10 Arabidopsis reference genome with the Araport11 annotation (89) using the HISAT2 software v2.1.0 (90). The alignment rate was about 99%. Transcript expression was quantified from uniquely mapped reads using the StringTie program v1.3.2 (91). StringTie’s prepDE.py script was used to extract read counts from the StringTie output. Fragments per kilobase of exon per million fragments mapped (FPKM) were calculated from StringTie output with the R package Ballgown (92). Pearson correlation coefficient was calculated between biological replicates with the normalised expression levels of log2 (FPKM+1). DESeq2 (93) was used for differential expression (DE) analysis. For exploring gene functions during analyses, the Araport11 annotation (89) .gff file was downloaded from arabidopsis.org. Custom R scripts (R v4.0.3) were used to parse the information and link it to the relevant genes during the analyses. To analyse sample variation, seed gene expression profiles were investigated using principal component analysis (PCA) and hierarchical clustering. For PCA, we first normalised the RNA-seq read counts of all genes with a variance stabilising transform (VST) using the R package ‘DESeq2’ V1.30.0 (93). We then conducted PCA using the R function prcomp with default settings and visualised the results using ggplot2.

### Dynamic time warping

To align time series data from round 2 and round 3 samples of Col-0 and *abi3-6*, dynamic time warping was performed using functions from the R package ‘dtw’ (v1.22.3) (94). The means of biological replicates were used for time warping to avoid assigning different time labels to individual replicates. Because phenotypic data from round 3 were more complete and used for correlations with both round 2 and round 3 expression data, Col-0 round 3 expression data were used as a template, while round 2 expression data were used as a query for alignment. The time axis alignment was configured with open beginnings and ends, and run across multiple step pattern constraints (’asymmetric’, ‘rigid’, and seven Rabiner–Juang step patterns), with the most frequently selected alignment taken as the final result. This alignment was applied to harmonize the mapping between round 2 expression time points and the corresponding phenotypic measurements from round 3 in Col-0. The majority of step patterns suggested a shift of approximately two days for early Col-0 round 2 samples (12–22 DAP), while later time points remained unchanged. For *abi3-6* round 2 samples, the same transformation was applied, as limited overlap with round 3 time points precluded a separate alignment.

### Gene co-expression network analysis

We constructed a co-expression network using the WGCNA package V1.69 (95) in R, including all 123 samples for the 11 available genotypes in the analysis. Three different criteria were used to filter the genes: having an expression of above 25 counts in at least 25% of the samples, at least 9 samples with an expression above 100 counts, or a coefficient of variation above 0.3. The expression of the remaining genes was normalized using the varianceStabilizingTransformation function from the R package DESeq2 V1.30.0 (93). We next checked for outliers among the samples by conducting hierarchical clustering using R’s hclust function with average linkage and Euclidean distance. For constructing the co-expression network calculated the adjacency of genes with a soft-thresholding power of 8 for a scale-free topology fit of a ‘signed hybrid’ network. The adjacency matrix was then transformed into a topological overlap matrix (TOM) to derive the weighted network. Genes were clustered hierarchically with average linkage, using 1-TOM as a distance metric. To separate the clustering result into modules, the function cutreeDynamic from the dynamicTreeCut package V1.63-1 was used with deepsplit=0 and a minimum required module size of 30 genes. Highly similar modules were merged with mergeCloseModules from the WGCNA package (Langfelder and Horvath, 2008), combining all modules with a dissimilarity of less than 0.14 to a final 17 modules with distinct patterns. We determined the functions associated with our WGCNA modules through Gene Ontology (GO) enrichment analysis of biological processes, which we conducted using topGO R package v2.42.0. To map GO terms to genes, we used the annotations available in the org.At.tair.db database. The significance of GO term enrichments in the modules was tested with the runTest function of topGO, selecting the ‘weight01’ algorithm and Fisher’s exact test to calculate p-values. All genes included in the WGCNA analysis served as the background for these tests. For multiple testing correction, we applied the BH-FDR correction method (96).

### Motif enrichment and transcription factor identification

Motif enrichment was performed with selected gene modules to find potential transcription factors (TFs) relevant to the traits. All genes from the selected modules were used for motif enrichment, except for the longevity-related modules turquoise and black because the large number of genes in these modules (>2000) would decrease the specificity of the analysis. Here, the EPPO data from all genotypes allowed for subsetting the genes based on ranked correlation. Genes falling in the top right quadrant of the module membership versus gene significance plot of the modules were selected to be used in motif enrichment (Supplemental Figure 1). The threshold gene significance (GS) and module membership (MM) were chosen to reduce the genes by about half, for the turquoise module: MM>0.78, GS>0.65, and black module: MM>0.8, GS>0.69. The genes were submitted to TF2Network (97)and the TFs found based on enriched position weight matrices were exported for further filtering. TFs of interest were manually selected based on their expression profiles for the different genotypes. Two selection criteria were applied. First, TFs showing an increase or decrease in expression in Col-0 at 18 DAP or 20 DAP were selected for longevity acquisition. Second, TFs with divergent expression profiles between mutants were prioritised, distinguishing genotypes that acquire longevity from those that do not. Some TFs could be further prioritised by showing that they are directly regulated by *ABI3* based on the research of (52), or by finding that they have genes as nearest neighbours that were previously shown in the literature to be important for the trait of interest, like *LEA*s for DT and longevity (11, 40, 42, 56, 57, 98). We considered genes nearest neighbours if they showed a topological overlap score of 0.2 or greater in the WGCNA TOM matrix.

### CRISPR-Cas9 mutants generation and phenotypic analyses

To generate CRISPR-Cas9 mutants, genes were selected based on the list of TFs related to longevity acquisition and filtered as described previously. To build the constructs, closely related genes (based on protein sequence) were targeted simultaneously in one construct (Supplemental Table 3). For each gene, we designed one or two guides using CHOPCHOP V3 (99) (Supplemental Figure 2, Supplemental Table 4). Target sites were selected for the lowest mismatch value and closest proximity to the translational start site, with a preference for the first exon, if specific enough. The guide-RNAs and subsequent expression vectors were cloned using the GoldenGate cloning system described by (100). As an expression vector, we chose the pDGE885 (pNOS:nptII_Ubq:FCY-UPP_pRPS5a:zCas9io_ccdB_CmR) to select transformants for kanamycin resistance and to later on select against transformants using the fungicide 5-fluorocytosine (5-FC) (100). Col-0 plants were transformed using floral dipping and T1 seeds were harvested in bulk for each construct and immediately germinated on ½ MS plates with 100 μg/ml kanamycin to select only successfully transformed seedlings. For each construct, 11 to 35 surviving seedlings were transferred to rockwool under greenhouse conditions and genotyped for signs of mutagenesis. Gene specific primers surrounding the target sites of each locus were used to genotype all lines (Supplemental Table 4). Primers were designed ∼150 bp upstream and ∼300 bp upstream of the target sites close to the ATG, and for genes targeted with two target sites an additional primer was designed ∼150 bp downstream of the distal target site. T1 seedlings showing a PCR product size differing from Col-0 were selected for further propagation. Seeds from each T1 parent were propagated separately on 1 mM 5-FC ½ MS plates to select against the Cas9 constructs. Eight surviving T2 seedlings per plate were transferred to rockwool for further selection and were genotyped as a pool for signs of mutagenesis. Pools of T2 plants that showed signs of mutagenesis were further genotyped by PCR as single plants. The individual T2 plants with mutations visible by PCR were selected for further propagation and, in addition, plants with a homozygous mutation were sequenced. Homozygous plants were then propagated and used for trial phenotyping experiments and T2 plants with one or more non-homozygous mutations were propagated for another round of selection for homozygosity in the next generation. For each CRISPR-Cas9 mutant line, four biological replicates, each consisting of four plants, were used, and seeds were harvested from each replicate independently. After three months of after-ripening (50% RH, 20°C), seeds were subjected to controlled deterioration treatment (CDT). For this, seeds were stored in the dark at 89% relative humidity (RH) and 40°C. Under these conditions, seeds remain in a dry state and do not germinate, distinguishing this treatment from heat stress applied to imbibed seeds. After 0, 2, 5, 7, 8, 9, and 10 days of storage, germination on dHLO was assessed. Maximum germination percentage (Gmax, %) was determined after seven days using Germinator software (80). For each replicate, Gmax values were fitted to a logistic curve to estimate the half-viability period (p50). Longevity differences between mutant lines and Col-0 were assessed by comparing p50 values using a Wilcoxon rank-sum test (α = 0.05).

## Results

### Physiological characterization of Arabidopsis seed maturation

Seeds from wild-type Columbia-0 (WT, Col-0) and of single gene mutants and mutant combinations of genes involved in ABA biosynthesis (*ABA2*), signalling (*ABI3, DOG1*), and catabolism (*CYP707A2*) (Supplemental Table 1) were harvested and phenotyped every two days from 12-14 to approximately 26 (dry seed, DS) days after pollination (DAP). We characterised temporal changes in the moisture content and acquisition of SQTs, including germination, dormancy and chlorophyll fluorescence, as well as DT and longevity in the DS stage (Figure 1A-F).

**Figure 1.**
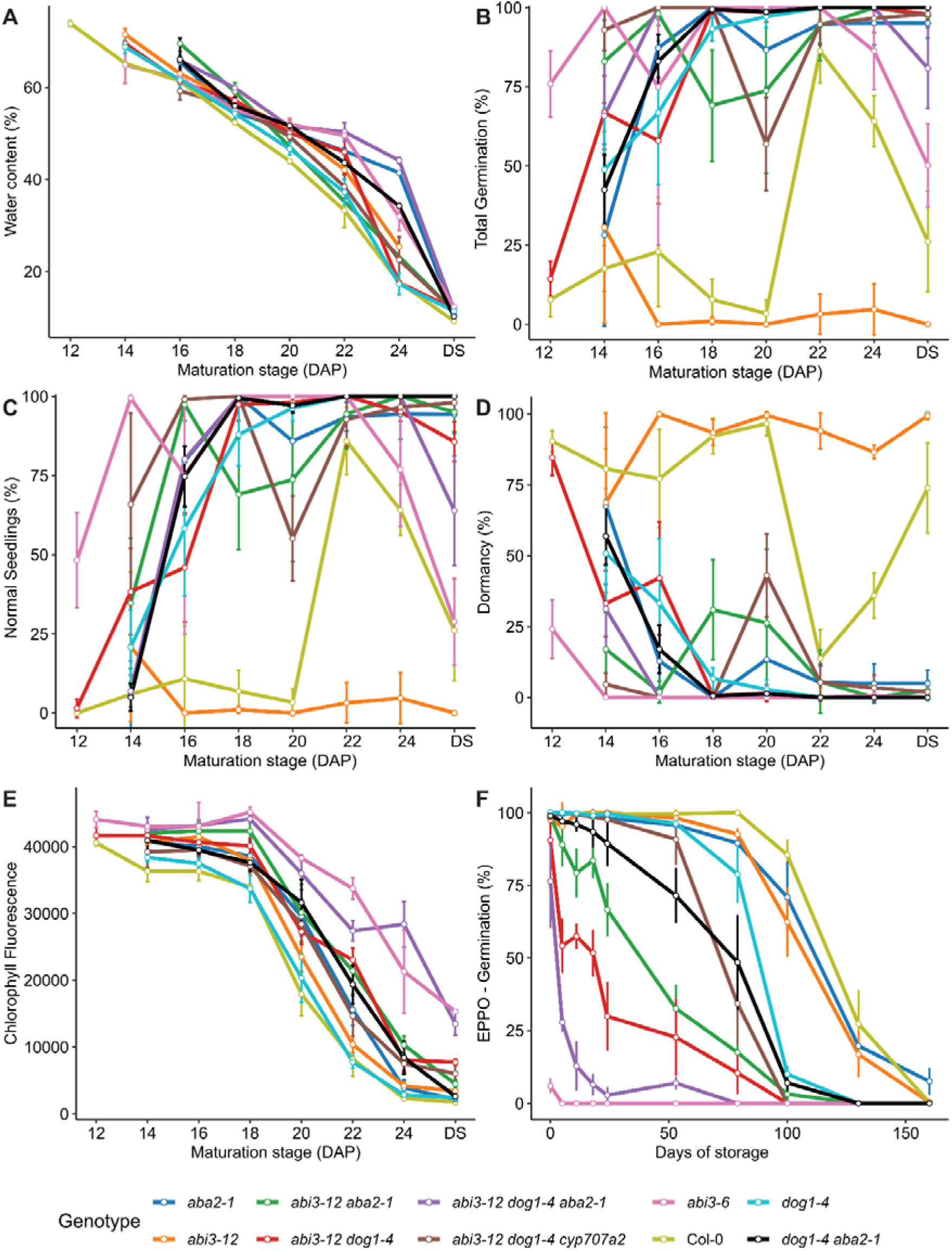
Seed maturation traits of the *Arabidopsis thaliana* wild type Col-0 and mutants *aba2-1, abi3-6, abi3-12, dog1-4*, *abi3-12 aba2-1, abi3-12 dog1-4*, *dog1-4 aba2-1*, *abi3-12 dog1-4 aba2-1* and *abi3-12 dog1-4 cyp707a2*. A - Water content percentage on a fresh weight basis, B - Total germination percentage (measured as radicle protrusion), C - Normal seedling percentage after germination (measured as seedlings with green cotyledons and elongated primary root), D - Dormancy percentage, E - Chlorophyll fluorescence (fv/fm), and F – Seed longevity measured as germination percentage after Elevated Partial Pressure of oxygen (EPPO) treatment. Each mutant genotype is represented by the same colours in all panels. A-E was performed using seeds from different maturation stages (days after pollination, DAP), n = 4. EPPO (panel F) treatment was performed in mature dry seeds (>26 DAP), n = 3.

The seeds of all mutants and WT reached around 10% water content at the end of seed maturation (DS stage, Figure 1A), with *aba2-1* and *ab3-12 dog1-4 aba2-1* mutants showing a remarkable slower decrease in moisture content at the end of maturation compared to Col-0. The ability to germinate (Figure 1B) and to produce normal seedlings (Figure 1C), as well as the primary dormancy level (Figure 1D) varied across the mutants throughout the maturation. For Col-0, the patterns of germination ability and dormancy were the opposite (Figure 1 B-D), with seeds showing low (10-20%) germination ability until 20 DAP, increasing to around 80% at 22 DAP and decreasing again to around 30% in DS. Except for *abi3-12*, all mutants showed overall higher germination and lower dormancy than Col-0 throughout the maturation (Figure 1B-D), with *abi3-6* showing high germination at the earliest maturation timepoint (12 DAP), while displaying low to no dormancy throughout the entire maturation time course (Figure 1B-D). In Col-0, a dramatic decrease in chlorophyll fluorescence was observed from 18 DAP onwards (Figure 1E). While chlorophyll fluorescence of all mutants also decreased during maturation; *abi3-6* and *abi3-12 dog1-4 aba2-1* showed a slower decrease in chlorophyll fluorescence when compared to Col-0. To assess longevity acquisition during seed maturation in a feasible manner, we performed controlled deterioration tests (CDT) in Col-0 at 2 and 5 days (Supplemental Figure 3A). Longevity increased from 20 DAP and peaked at 22 DAP under 2-day CDT, whereas under 5-day CDT acquisition was delayed, increasing from 22 DAP and peaking at the DS stage. To compare longevity between genotypes, we used the elevated partial pressure of oxygen (EPPO) assay on seeds at the DS stage. All mutants showed reduced longevity compared to Col-0 upon EPPO (Figure 1F). The low germination of *abi3-6* even without EPPO (day 0) reflects its well-known desiccation-sensitive phenotype. During storage, Col-0 longevity began to decline after ∼80 days of EPPO, followed by *abi3-12*, *aba2-1*, and *dog1-4*, while stronger reductions were observed in double and triple mutants combining *abi3-12* with defects in ABA biosynthesis or signalling. To also assess DT acquisition during seed maturation, we subjected maturing Col-0 seeds at multiple developmental stages to drying. This analysis showed that DT increases between 12 DAP and 16 DAP (Supplemental Figure 3B). Since DT is defined as the ability to survive near-complete water loss and resume metabolism upon rehydration (101), seeds of all mutants, except *abi3-6*, were able to acquire DT, as indicated by seeds at the DS stage showing low moisture content (∼10%) (Figure 1A) and being able to germinate prior to EPPO storage (day 0, Figure 1F). These observations suggest that, although DT is ultimately achieved in most mutants, its temporal regulation during maturation may be altered, warranting further investigation.

To understand the contribution of ABA to SQT acquisition during maturation, we quantified ABA at 18 DAP and in dry seeds of Col-0 and all mutants (Supplemental Figure 4A-B). As expected, the single and combinations of the *aba2-1* mutant (*abi3-12 aba2-1* and *abi3-12 dog1-4 aba2-1*) had significantly lower ABA when compared to Col-0 at 18 DAP, indicating that decreased *ABA2* expression is sufficient to decrease the levels of ABA during seed maturation in these mutants (Supplemental Figure 4A). On the other hand, *abi3-12* and *abi3-12 dog1-4 cyp707a2* showed significantly higher ABA content. The higher accumulation of ABA in *abi3-12* could explain why this mutant displays much higher dormancy than all other mutants and the Col-0 during seed maturation (Figure 1D), and loss of ABA catabolism enzyme CYP707a2 in the *abi3-12 dog1-4 cyp707a2* mutant explains the higher ABA content in seeds at 18 DAP. At the DS stage all mutants had similar seed ABA content as the wild-type, except for *abi3-12 dog1-4 cyp707a2* triple mutant which had significantly higher ABA content (Supplemental Figure 4B) suggesting that, as long as *CYP707a2* is fully functional, seeds will accumulate wild-type levels of ABA at the end of maturation.

Together, these results reveal that SQT acquisition during maturation is highly dynamic and differentially regulated across genotypes, highlighting the complex and combinatorial nature of seed maturation regulation.

### The transcriptome landscape of Arabidopsis seed maturation

To gain molecular insight into this dynamic regulation of SQTs, we systematically identified transcriptional signatures associated with Arabidopsis seed maturation. We performed RNA sequencing (RNA-seq) of Col-0 seeds across eight stages, every two days from 12 DAP to the dry seed stage (∼26 DAP, DS) (Supplemental Table 2). In addition, RNA-seq was performed on the seed maturation mutants at selected time points chosen to capture stages most relevant to specific SQTs. Using principal component analysis (PCA) a clear separation between the different genotypes was observed in the first principal component (PC1), explaining 40.4% of the total variability, while PC2 shows a clear progression of the maturation time for each genotype, contributing with 31.52% of the variability (Figure 2A).

**Figure 2.**
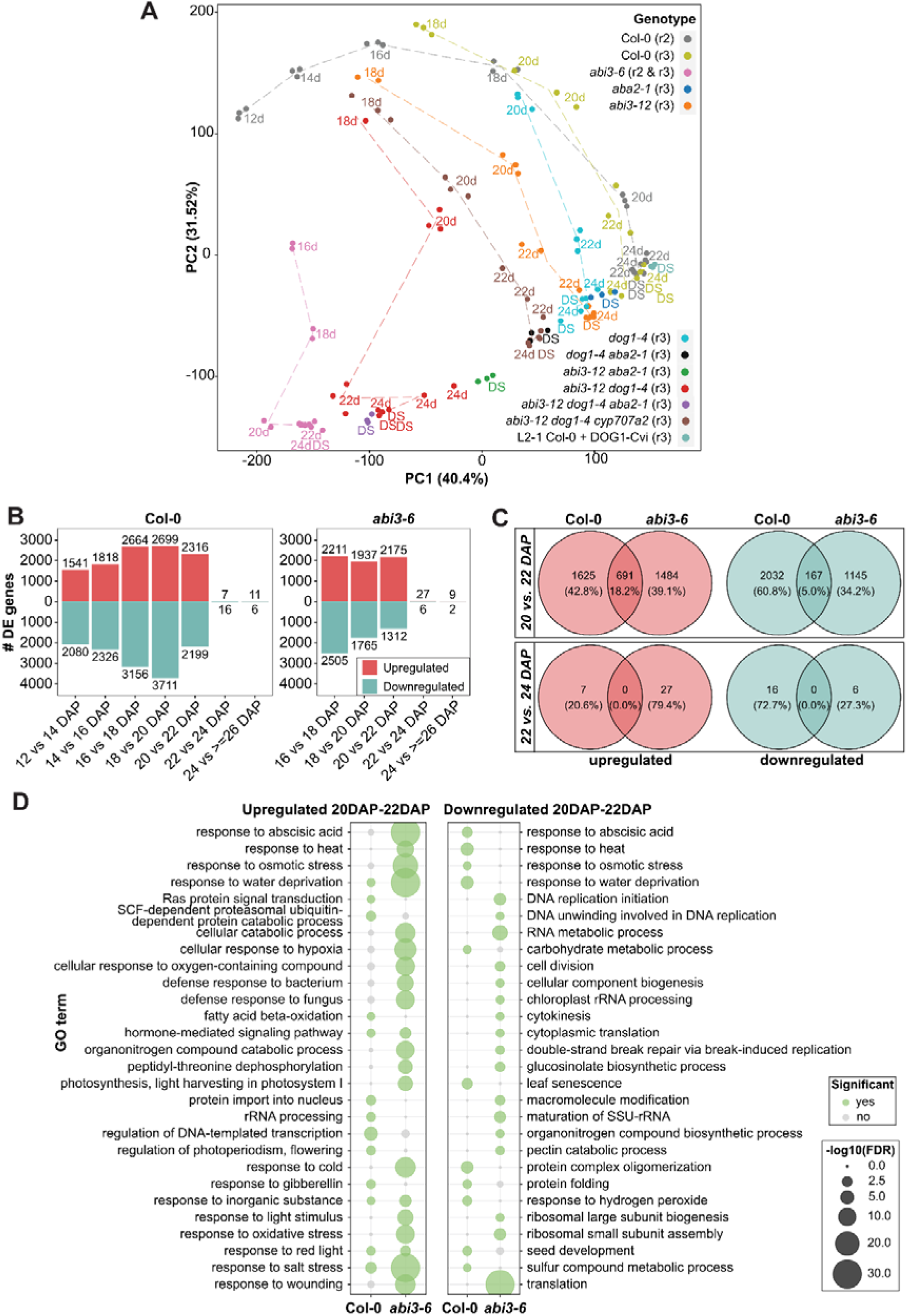
Transcriptome landscape of *Arabidopsis thaliana* seed maturation. A - Principal component analysis (PCA) of variance stabilizing transformation-normalized RNA-seq read counts showing global transcriptome relationships across different days (d) of seed maturation and dry seeds (DS) in mutants and wild-type (Col-0) (123 seed samples). Experimental rounds 2 (r2) and 3 (r3) are indicated between parenthesis (Supplemental Table 2). B - Number of differentially expressed (DE) genes of Col-0 (round 2) and *abi3-6* (round 2 and 3) between consecutive days after pollination (DAP). C - Numbers of shared and unique up and downregulated genes (p-adj. < 0.01) for Col-0 and *abi3-6* seed maturation between 22 DAP and 20 DAP, and between 24 DAP and 22 DAP. D - Gene ontology (GO) term enrichment of shared and uniquely upregulated and downregulated genes in Col-0 and *abi3-6* between 20 DAP to 22 DAP (p-adj. < 0.05). The top 20 representative enriched GO categories among the upregulated (left) and downregulated (right) genes are shown. The green colour indicates significance, and size of the dots represent enrichment level (-log_10_p-adj.).

We observed a gradual separation in the maturation time course among the various mutants compared to the maturation samples of Col-0. The *dog1-4* and *abi3-12* had a least extreme transcriptome profile change compared to Col-0 for their respective maturation time course samples, and all genotypes containing the *abi3-12* mutation (*abi3-12 aba2-1*, *abi3-12 dog1-4*, *abi3-12 dog1-4 aba2-1 and abi3-12 dog1-4 cyp707a2*) showed greater separation between their samples compared to the same time points in Col-0 (Figure 2A). Remarkably, the *abi3-6* maturation time course was notably the most distinct from Col-0, indicating major transcriptional changes during seed maturation upon severe *ABI3* impairment. To initially explore our datasets and to investigate how gene expression dynamics during maturation are altered between contrasting genotypes, we performed differential expression (DE) analysis between consecutive timepoints in Col-0 and in *abi3-6* (Figure 2B). In Col-0, the number of up and downregulated genes between consecutive time points increased until 20 DAP (Figure 2B, Supplemental Data Set 1). From 22 DAP to 24 DAP the striking decrease in both up and downregulated genes indicates a global transcriptome shutdown as the seeds reach the dry state. In *abi3-6*, the number of upregulated genes remained relatively similar until 22 DAP, while the number of downregulated genes gradually decreased. From 22 DAP to 24 DAP a striking decrease in both up and downregulated genes, similar to Col-0, was observed (Figure 2B, Supplemental Data Set 2). To identify transcriptional signatures that could explain the contrasting maturation phenotypes of Col-0 and *abi3-6*, we compared genes that were up- and downregulated preceding or after the global transcriptome shutdown in both genotypes (Figure 2C). A small percentage of upregulated (18.2%) and downregulated (5.0%) genes were shared between Col-0 and *abi3-6* before the transcriptome shutdown (20 - 22 DAP), and no up- or downregulated genes were shared between both genotypes after the transcriptome shutdown (22 - 24 DAP) (Figure 2C). This suggests that, despite a similar transcriptome shutdown before reaching the dry seed stage, Col-0 and *abi3-6* have very distinct transcriptome profiles. Gene Ontology (GO) enrichment analysis revealed distinct enrichment of biological processes between Col-0 and *abi3-6* before the transcriptome shutdown (20 - 22 DAP) (Figure 2D, Supplemental Data Set 3). Overall, *abi3-6* showed upregulation of processes related to stress response, such as response to abscisic acid, response to osmotic stress, response to water deprivation, cellular response to hypoxia, response to cold, and response to salt stress, several of these stress-related processes were significantly downregulated in Col-0. In contrast, *abi3-6* showed significant downregulation of processes such as translation and RNA metabolism which were not downregulated in Col-0. Together, these observations indicate that *abi3-6* exhibits a stress-associated transcriptional signature distinct from Col-0, along with a reduced capacity to properly coordinate transcriptional programs required for seed maturation and DT.

To facilitate exploration and integration of these datasets, we generated a user-friendly web interface, SeedMatExplorer (https://www.bioinformatics.nl/SeedMatExplorer), which enables comprehensive visualization of the phenotyping data alongside gene expression analyses across mutants. The platform further provides access to differential expression (DE) analyses, gene ontology (GO) enrichment, and weighted correlation network (WGCNA) analyses (described below), allowing users to interrogate transcriptomic patterns underlying seed maturation in a flexible and interactive manner.

### Multiple co-expressed gene modules are associated with specific SQTs

Because the transcriptome data for Col-0 and *abi3-6* were generated in two independent experimental rounds, we leveraged this design to robustly align the RNA-seq time-series using dynamic time warping (DTW) (Giorgino, 2009), thereby enabling the integration of all 123 RNA-seq samples across mutants and timepoints (Supplemental Table 2) for weighted correlation network analysis (WGCNA). This approach facilitated a comprehensive network-level analysis, identifying 17 gene modules with distinct expression patterns, summarized as module eigengenes (MEs), representing the weighted average expression profiles across genotypes and developmental stages (Figure 3A, Supplemental Data Set 4). The effect of dynamic time warping is illustrated by comparing WGCNA module eigengenes with and without time warping of round 2 samples for Col-0 and *abi3-6* (Supplemental Figure 5A–B).

**Figure 3.**
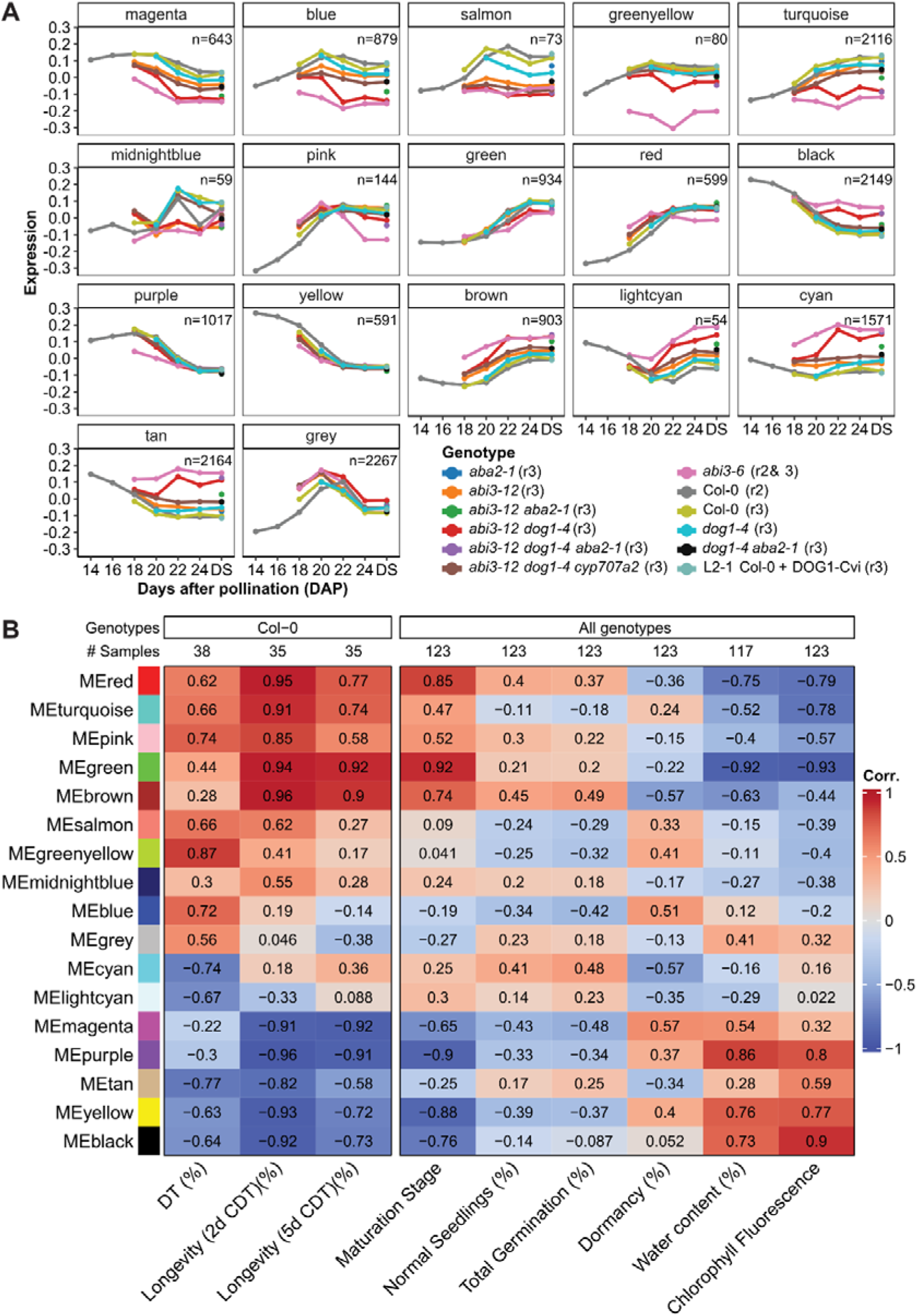
Module eigengenes (MEs) generated by WGCNA. A - ME expression profiles, representing the expression of the module’s genes for each genotype throughout the maturation time course. These profiles are the first principal component of expression read counts that have been normalised with variance stabilising transformation. The numbers in the upper right corner indicate gene count in individual clusters. DS = Dry Seed. B - Module eigengene (ME) associations with the indicated seed physiological traits. The scale on the right indicates the Pearson correlation coefficient (PCC). The numbers at the top indicate the number of samples used for the analysis. DT = desiccation tolerance, CDT = controlled deterioration test.

The largest modules contained around 2200 genes (grey, tan, and black modules) and the smallest module contained 54 genes (lightcyan) (Figure 3A). Interestingly, the expression pattern of some modules, for example magenta, blue, turquoise, and cyan, clearly followed a similar genotype separation pattern as we have observed in the PCA (Figure 2A), potentially containing genes underlying transcriptional and phenotypic differences between the different genotypes. The greenyellow module showed a strikingly different expression profile in *abi3-6* compared to the other genotypes, suggesting that it may contain genes underlying the severe maturation defect of *abi3-6* (Figure 1B-F).

To further explore the gene modules and identify genes and transcription factors (TFs) associated with SQT regulation, we used Pearson’s correlation coefficients (PCCs) to evaluate correlations between the 17 WGCNA modules and the available trait data for each genotype (Figure 3B). Multiple distinct modules exhibited both positive and negative correlations with the same traits, suggesting that SQT regulation is governed by multiple gene groups with distinct expression patterns. For Col-0 SQTs, DT acquisition showed the highest positive correlation with the greenyellow module (PCC = 0.87) and the strongest negative correlation with the tan module (PCC = -0.77). In the longevity datasets (2 d and 5 d CDT, Supplemental Figure 3), multiple strong correlations were observed, including high positive correlations (PCC > 0.9) with the brown, red, green, and turquoise modules, and strong negative correlations (PCC < −0.9) with the purple, yellow, magenta, and black modules. Using phenotypic data from all genotypes, we also identified modules associated with additional seed traits. Seed maturation stage (DAP) showed strong positive correlations with the green (PCC = 0.92) and red (PCC = 0.85) modules, and strong negative correlations with the purple, yellow, and black modules. An opposite correlation pattern was observed for water content and chlorophyll fluorescence, consistent with these traits reflecting maturation stage. In contrast, correlations between gene modules and the remaining traits (normal seedlings, germination, and dormancy) were weaker, ranging between −0.57 and 0.51.

### Novel gene modules associated with desiccation tolerance and longevity in Arabidopsis

Given the importance of desiccation tolerance (DT) and longevity for seed resilience and survival in dry environments and during storage, and the complex and still incompletely understood regulation of these traits, we focused on the modules most relevant to these processes. To systematically explore their biological relevance and identify potential regulators, we integrated Pearson correlation analysis, Gene Ontology (GO) enrichment, and enrichment of *LEA* genes. This combined approach enabled us to link transcriptional modules to specific seed traits and functional categories. Initially, we evaluated the Pearson correlation between module eigengenes and the DT and longevity trait values. Because Col-0 provided the most complete time-series trait data (Supplemental Table 2), correlation analysis was performed using this genotype. Modules showing a high correlation (PCC > 0.8) with DT or longevity in Col-0 were considered as candidates for further investigation (Figure 3B). To avoid bias based on expression-trait associations in Col-0 alone, we did not select modules solely on the basis of correlation strength. Instead, we used the direction and relative differences in module eigengene profiles across genotypes (e.g., differences between genotypes with higher versus lower longevity, Figure 1) as a qualitative consistency check to ensure that modules associated with a given trait in Col-0 did not show conflicting expression patterns across genotypes. This step was used to support the biological relevance of candidate modules rather than to exclude modules based on arbitrary criteria. To further refine module prioritization and address the limitations of a correlation-based approach, namely that gene expression changes may precede measurable trait differences and that correlations were derived from Col-0 only, we applied additional, independent lines of evidence. In particular, modules approaching or exceeding the correlation threshold were further supported if they showed enrichment for Gene Ontology (GO) terms related to abiotic stress responses (Supplemental Figure 6) and/or enrichment for *LEA* genes, assessed using Fisher’s exact test with the full WGCNA gene set as background (Supplemental Table 5). *LEA* enrichment was considered significant at p < 0.05.

Among the WGCNA modules, the greenyellow module showed the highest correlation with DT acquisition and exhibited markedly lower expression in *abi3-6*, consistent with the desiccation-sensitive phenotype of this mutant (Figure 3A). Given the established role of *ABI3* as a master regulator of DT, we examined the extent to which genes in this module are associated with *ABI3* regulation. Of the 80 genes in the greenyellow module, 66 (82.5%) have been reported as ABI3-associated based on ChIP-seq or transcriptome studies, including 52 genes (65%) directly and positively regulated by ABI3 (51, 52, 73); Supplemental Table 6 and 7). Notably, the greenyellow module showed the highest enrichment of *LEA* genes among all modules, with 11 genes (13.8%), of which 9 are direct ABI3 targets (Supplemental Table 5; (52)). Together with its strong correlation with DT acquisition, enrichment of ABI3-regulated genes (Figure 3B, Supplemental Table 7), and the well-established association of LEAs with DT (98, 102, 103), this module represents a strong candidate for containing genes involved in DT acquisition in Arabidopsis seeds. Comparison with previously identified DT-associated co-expression modules (73) showed that 60 genes (75%) overlap with DT-associated genes, while 5 genes were not reported in those datasets (Figure 4A, Supplemental Table 8). These genes encode proteins with predicted or reported roles in RNA modification, small RNA regulation, and DNA repair (Figure 4B) (104–108). While these genes have not been functionally characterized in the context of DT, their co-expression with known *ABI3*-and DT-associated genes highlights them as candidates for further investigation.

**Figure 4.**
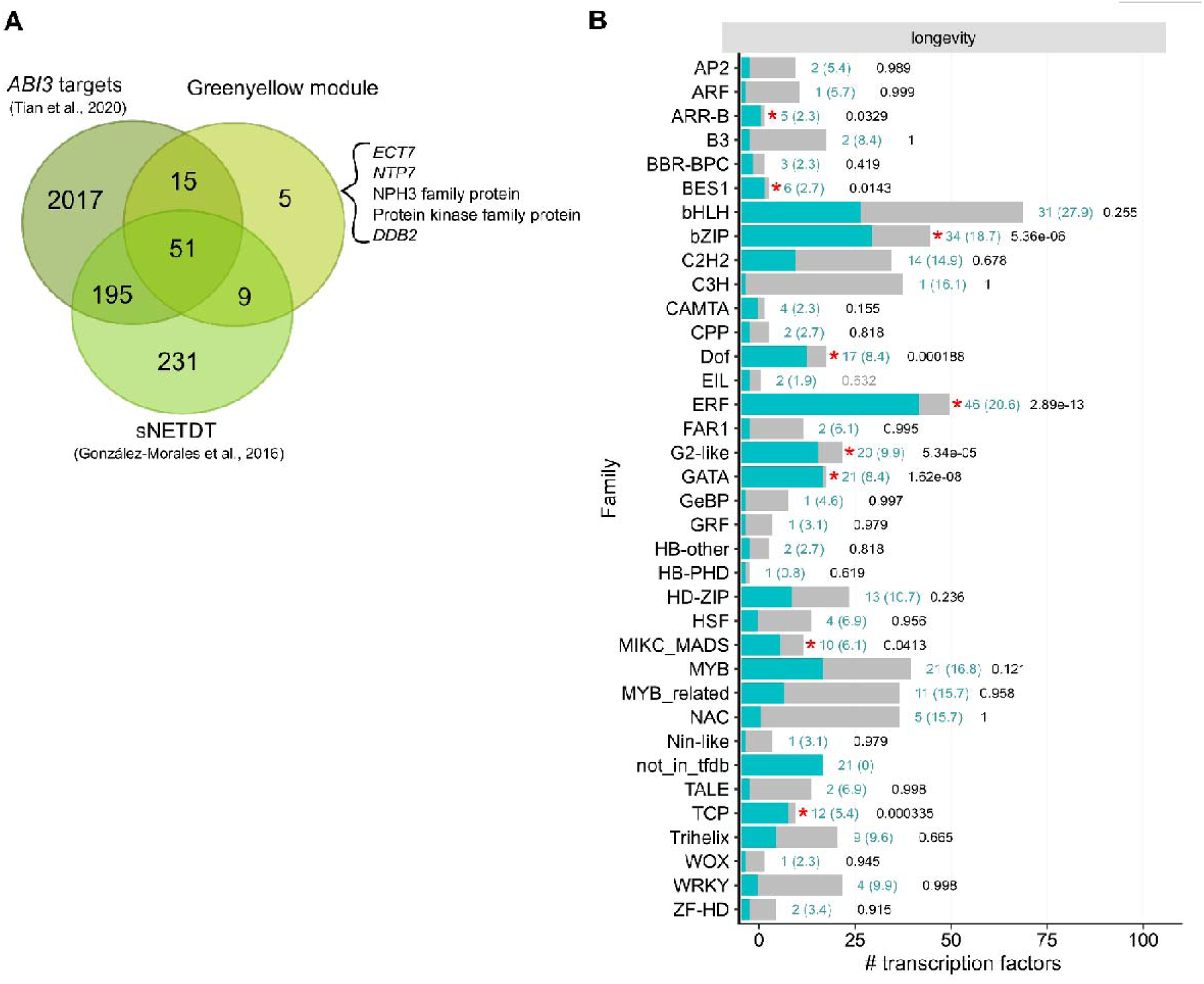
Identification of novel genes and gene modules associated with DT and longevity. A - Venn diagram of genes identified as direct or indirect targets of ABI3 through ChIP-seq and transcriptome analysis (Tian et al., 2020), genes from two Arabidopsis seed DT co-expression subnetworks (sNETDT, González-Morales et al. (2016), and genes from the greenyellow module. Genes uniquely found in the greenyellow module are indicated after braces. B - In turquoise: TF families identified by motif enrichment analysis with TF2Network for longevity modules. TF families were determined based on the plantTFDB list; the background distribution is shown as grey bars. The numbers adjacent to the bars indicate the number of TFs that were found in TF2Network analysis, and parentheses show the number of TFs that would have been expected by chance. Black values show the BH adjusted p-values (Benjamini and Hochberg, 1995) for Fisher’s exact test. The result includes all transcription factors from TF2Network analysis that passed filtering for low expression and low variation, using the same thresholds as in the WGCNA analysis (see Materials & Methods). Red asterisks indicate significant enrichment of binding motifs in the dataset (p<0.05).

Longevity is a complex quantitative trait influenced by genetic factors and environmental conditions during seed maturation and storage (11). Consistent with this, multiple gene modules showed strong correlations with longevity acquisition in Col-0 based on CDT data (Figure 3B), including positive correlations for the brown (PCC = 0.96), red (PCC = 0.95), green (PCC = 0.94), and turquoise (PCC = 0.91) modules, and negative correlations for the magenta (PCC = -0.92), purple (PCC = -0.96), yellow (PCC = -0.93), and black (PCC = - 0.92) modules. To identify potential regulators of these longevity-associated modules, we performed transcription factor (TF) enrichment analysis using TF2Network (Kulkarni et al., 2018), focusing on expressed TFs present in the WGCNA dataset. This yielded 332 candidate TFs, with enrichment of several TF families, including ARR-B, BES1, bZIP, Dof, ERF, G2-like, GATA, MIKC_MADS, and TCP (Figure 3C; Supplemental Data Set 5), suggesting coordinated transcriptional regulation underlying seed longevity. Using multiple filtering steps (see Materials & Methods), we identified 25 TFs as top candidates potentially associated with regulation of longevity acquisition during seed maturation (Supplemental Table 9). Among these, several belong to the DREB and EAR motif protein (DEAR) family within the ERF/AP2 DREB subfamily A-5, including DEAR4 (RAP2.10), DEAR5 (RAP2.9) and RAP2.1, which have been described as stress-inducible transcriptional repressors (109–112). As an initial screening and exploratory validation of the role of these candidate TFs in seed longevity, we generated single and higher-order CRISPR-Ca9 mutants of these tree TFs and the closely related DEAR2 TF (Supplemental Table 3-4, Supplemental Figure 2) and we assessed their germination after storage under natural conditions and after CDT. Some combinations, such as *dear2rap2.1*, *dear2dear4rap2.1* and *dear2dear4dear5rap2.1*, showed reduced germination compared to Col-0 after natural storage (Supplemental Figure 7A). Under CDT, *dear2rap2.1* showed significantly decreased viability while *dear5* and *dear4dear5* showed significantly increased viability compared to Col-0, as shown by the higher and lower number of days to reduce 50% viability, respectively (*p*50, Supplemental Figure 7B). These observations support the identification of gene modules with novel candidate regulators of seed longevity, and highlight TFs for further functional investigation.

## Discussion

We integrated extensive physiological analyses with RNA-seq data of wild-type (Col-0) and mutants affected in key regulators of seed maturation, including *ABI3* and ABA biosynthesis and catabolism genes, to identify groups of genes associated with the acquisition of distinct seed quality traits (SQTs). Although most mutants studied here ultimately reached similar low moisture content and acquired DT by the dry seed stage, several showed delayed maturation-associated processes, including slower water loss and chlorophyll degradation, as well as altered dormancy dynamics (Figure 1). Notably, while most mutants displayed reduced dormancy and increased germination during maturation, *ABI3* mutants exhibited more complex phenotypes, including higher dormancy of *abi3-12* and lower longevity and desiccation sensitivity of *abi3-6*, in agreement with the established role of ABI3 as a central regulator of seed maturation, dormancy, and DT (34, 38). The longevity assays further revealed that, despite successful DT acquisition, all mutants (except *abi3-6*) were compromised in germination after EPPO storage, particularly higher-order combinations. These observations suggest that DT and longevity are mechanistically distinct and can be genetically uncoupled, as previously reported (7, 28, 34, 35, 56, 57, 59, 64, 73). ABA measurements showed that perturbations in ABA metabolism significantly affected dormancy and maturation timing, but had limited impact on final ABA levels in dry seeds, unless ABA catabolism was impaired. Overall, our findings highlight the dynamic and combinatorial role of ABA-related pathways in fine-tuning SQTs during maturation.

We then generated a comprehensive gene expression atlas of Col-0 and of ABA biosynthesis, catabolism, and signalling mutants to study the underlying transcriptional control of SQT acquisition (Supplemental Table 1). *ABI3* mutation strongly affects overall seed development, especially the accumulation of seed storage and LEA proteins (70, 113), chlorophyll degradation (41); DT and longevity acquisition (34) and dormancy acquisition (30). In line with this, our RNA-seq data analysis revealed that mutation on the strong allele of *ABI3* (*abi3-6*) leads to an extreme transcriptome shift during seed maturation when compared to Col-0 (Figure 2A), although both genotypes showed a transcriptome shutdown from 22 DAP onwards after the onset of drying (Figure 2B, Figure 1A). We found a significant upregulation of stress-related biological processes during *abi3-6* maturation contrary to Col-0, just before the global transcriptome shutdown at 22 DAP (Figure 2C). *ABI3* mutation has been shown to directly affect the expression of stress responsive genes in seeds under osmotic stress (114) and in seedlings under dehydration stress (Bedi et al., 2016). It is possible that, with seed drying at around 20 DAP, the functional impairment of *ABI3* function in *abi3-6* leads to an enhanced expression of stress-responsive genes instead of the expected expression of DT-related genes, such as *LEA*s (51, 52, 73).

By combined analysis of the RNA-seq data and the phenotypic data of SQT acquisition for Col-0 and ABA-related mutants, we identified seventeen distinct gene expression modules which were correlated with specific SQTs (Figure 3). The acquisition of DT, acquisition of longevity, and the maturation stage have a relatively similar module-trait correlation pattern (Figure 3B), indicating that, although DT and longevity are temporally uncoupled, they might be similarly influenced by the seed developmental stage. On the other hand, the correlation similarity between seed moisture and chlorophyll fluorescence was the opposite of what we observed for DT acquisition, longevity, and maturation stage. Although the physiological relationship between seed moisture content decrease and chlorophyll content has been previously described for crop species (115–118), the effects of seed drying on chlorophyll degradation during seed maturation have not yet been explored at the molecular level, representing an interesting physiological aspect of seed maturation for future investigation.

An in-depth analysis of the gene module with the highest correlation with DT (greenyellow module) revealed a strong enrichment of genes previously described as part of the ABI3 regulon and associated with DT (52, 73) (Figure 4A). Among these, we identified a subset of genes that, to our knowledge, have not yet been linked to DT and may represent candidates for further investigation. Longevity is a complex quantitative trait regulated by the combinatorial action of multiple TFs (11, 119, 120). Our analyses identified four gene modules strongly positively correlated with longevity (brown, red, green, and turquoise; Figure 3B), along with enriched promoter TF-binding motifs and candidate TFs associated with seed longevity (Figure 4). Among these, DEAR TFs emerged as particularly promising candidates for seed longevity.

## Conclusions

Our study provides a comprehensive and integrated resource that advances the understanding of regulatory mechanisms underlying seed maturation in *Arabidopsis thaliana*, particularly the acquisition of SQTs essential for survival, resilience, and storage. By combining high-resolution transcriptomic data with detailed physiological characterization across wild-type and ABA-related mutants, we identified gene modules and regulatory programs associated with the temporal acquisition of key traits, including DT and longevity. These findings highlight the complex and coordinated nature of gene regulation during seed maturation. To facilitate data accessibility and reuse, we developed SeedMatExplorer (https://www.bioinformatics.nl/SeedMatExplorer), a user-friendly platform that enables the community to explore these datasets and generate new hypotheses on the transcriptional regulation of seed maturation and associated traits. Together, this resource provides a valuable framework for future studies on seed biology and has potential applications in improving seed performance and storage in agricultural systems.

## Ethics approval and consent to participate

Not applicable.

## Consent for publication

Not applicable.

## Availability of data

The sequencing data generated in this study are available in the Gene Expression Omnibus (GEO) under accession number GSE270988. Processed data can be found in the SeedMatExplorer browser https://www.bioinformatics.nl/SeedMatExplorer/app/. The plant material used and generated during the current study are available from the corresponding author on reasonable request.

## Author contributions

B.J.W.D., A.C.S.S.M. and R.N.T. designed the research.

M.A.S.A., L.A.J.W., L.L.B., A.D.D., J.A.D., B.J.W.D., A.C.S.S.M.., A.A.S. and R.N.T. performed seed physiological experiments.

L.L.B., A.D.D., J.A.D. and R.N.T. designed and generated CRISPR mutants. R.A.K., H.N. and R.N.T. performed bioinformatics analysis.

M.A.S.A., R.A.K., L.L.B., A.A.S. and R.N.T. analysed experimental data. M.A.S.A., R.A.K. and R.N.T. wrote the manuscript.

B.J.W.D., L.B., H.H. and R.N.T. conceived and supervised the project.

All authors commented on the article and agreed upon the final version of the manuscript.

## Acknowledgements

We thank all members of the Wageningen Seed Science (WSSC) for their contribution with the experiments, Francel Verstappen from the Laboratory of Plant Physiology of Wageningen University and Research for his help with hormone measurements and Dr. Parvinderdeep Kahlon from Laboratory of the Plant Physiology from Wageningen University and Research for his insightful feedback on the manuscript.

## Supplementary Information

### Additional file 1 – Supplemental Figures

**Supplemental Figure 1.** Subsetting of the genes from WGCNA modules ‘black’ and ‘turquoise’ to select those most relevant to longevity acquisition. The genes are separated on the scatter plots’ x-axis based on module membership (MM), the correlation of a gene’s expression profile with the module eigengene of its respective WGCNA module, and on the y-axis by the correlation of their dry stage expression across genotypes with longevity ranks for those genotypes from the EPPO experiment. Only genes that fall in the area shaded green (top-right) were included in longevity motif enrichment analysis.

**Supplemental Figure 2.** Overview of CRISPR-Cas9 mutant construct generation. Sg1 - Single guide RNA 1.

**Supplemental Figure 3.** Physiological characterization of longevity (A) and desiccation tolerance (DT) (B) acquisition during Col-0 seed maturation. A - Controlled deterioration test (CDT) was performed on maturing seeds every two days from 16 days after pollination (DAP) up to the dry seed (DS) stage. Seeds were stored for 2 days or 5 days at 80−85 % relative humidity (RH) and 40 °C prior to germination (n = 4). B – DT assessment was performed on maturing seeds every two days from 12 days after pollination (DAP) up to the dry seed (DS) stage. Seeds were desiccated for 48 hours at 22 °C and 30% relative humidity (RH) in the dark prior to germination (n = 4). Control consisted of seeds not desiccated and directly submitted to germination. DT percentage was calculated as the difference in the percentage of normal seedlings after 7 days between desiccated and non-desiccated samples. For both A-B, germination was performed on 10 μM GA_4+7_ and 10 mM KNO_3_ at 22 °C under continuous light.

**Supplemental Figure 4.** ABA content in (A) 18 DAP and (B) dry seeds. Values are normalised by 5 mg of seeds and internal standard (IS) by LC-MS/MS, (n = 4). ANOVA was conducted to test for differences in mean ABA content across genotypes, pairwise comparisons were made using Tukey’s HSD post-hoc test. In the figure, a compact letter display illustrates statistically similar means: genotypes sharing the same letter in panel A or B have means that are not significantly different. Specifically, a shared letter indicates a Tukey HSD p-value above the significance threshold of 0.05.

**Supplemental Figure 5.** Effect of dynamic time warping on module eigengenes (MEs). A. MEs without warping time axis. B. MEs with warped time axis. Time warping was used to adjust time point labels of round 2 to round 3, only Col-0 and *abi3-6* samples are affected. Adjustment of time axis labels for round 2 samples was as follows: +2 days for each sample from 12 to 22 days. This figure shows a better agreement of the ME profiles of Col-0 round 2 (grey) and Col-0 round 3 (lightgreen), clearly noticeable when comparing for the black module, before and after time-warping. Improved alignment of time-series for abi3-6 round 2 samples is noticeable when comparing the ME of *abi3-6* (pink) with the ME of *abi3-12 dog1-4 (red)*, for example for the turquoise module, before and after time-warping.

**Supplemental Figure 6.** Overview of significantly enriched GO terms and trait correlations for modules obtained from WGCNA analysis. The selection of GO terms includes all biological processes with a significant enrichment (padj <0.05). A column with grey text displays the number of genes in each module annotated with a GO term, as well as the total number of genes annotated with the term in the background population. The expected number of genes associated with each GO term is shown based on the total number of genes in the module and the prevalence of the term in the background population. At the bottom of each graph, a heatmap is presented depicting the Pearson correlation between the eigengene of each module and a set of traits. The p−value corresponding to the correlation shown in parentheses.

**Supplemental Figure 7.** A - Percentage of germination of mature dry seeds stored under natural conditions (50%RH, 20°C) for 47 days from Col-0 and multiple CRISPR-Cas9 mutants germinated in the presence of 10 μM GA_4+7_ and 10 mM KNO_3_. B - Half viability time (number of days to lose 50% of seed viability) of seeds submitted to CDT. Asterisks in A and B indicate significant differences (p<0.05) based on a Wilcoxon Sum Rank test and error bars indicate standard error (n = 4). The different CRISPR alleles and backgrounds are described in Supplemental Figure 3.

## Additional Files 2 - Supplemental Tables

**Supplemental Table 1**. Overview of genotypes used in this study and their respective phenotypes.

**Supplemental Table 2**. RNA-seq sample overview. Number of samples sequenced per time point (DAP = days after pollination, DS = Dry Seed).

**Supplemental Table 3**. Summary of CRISPR-Cas9 lines generated in this study.

**Supplemental Table 4**. List of guide RNAs and genotyping primers used for CRISPR-Cas9 mutants generated in this study.

**Supplemental Table 5**. Enrichment of LEAs amongst the ABI3 regulated genes for WGCNA modules. P-values were obtained with Fisher’s exact test.

**Supplemental Table 6**. Description and ABI3 regulation of genes belonging to the greenyellow module (DT). LEA genes are indicated in bold.

**Supplemental Table 7**. Enrichment of ABI3 regulated genes for WGCNA modules. P-values were obtained with Fisher’s exact test.

**Supplemental Table 8**. Genes that are direct or indirect targets of ABI3 based on Tian et al. (2020), genes from the greenyellow module (this work) and genes of two desiccation-tolerance subnetworks based on González-Moralez et al. (2016).

**Supplemental Table 9**. Filtered lists of candidate transcription factors involved in longevity regulation in Arabidopsis. Genes selected for CRISPR-Cas9 mutant validation are indicated in bold.

## Additional File 3 - Supplemental Data Sets

**Supplemental Data Set 1.** Differentially expressed (DE) genes between individual subsequent days after pollination (DAP) for Col-0. AGI – gene identifier, log2FC – log2 fold-change, stat - statistics, p-adj – false discovery rate adjusted p-value <0.01.

**Supplemental Data Set 2.** Differentially expressed (DE) genes between individual subsequent days after pollination (DAP) for abi3-6. AGI – gene identifier, log2FC – log2 fold- change, stat - statistics, p-adj – false discovery rate adjusted p-value <0.01.

**Supplemental Data Set 3.** Gene Ontology (GO) term enrichment for downregulated and upregulated differentially expressed (DE) genes at 20 and 22 days after pollination (DAP)

**Supplemental Data Set 4.** List of TFs identified using TF2Network and in the WGNA analysis.

**Supplemental Data Set 5.** Genes belonging to 17 modules created with WGCNA.

