## Supplemental Figures for "SeedMatExplorer: The transcriptome atlas of Arabidopsis seed maturation"

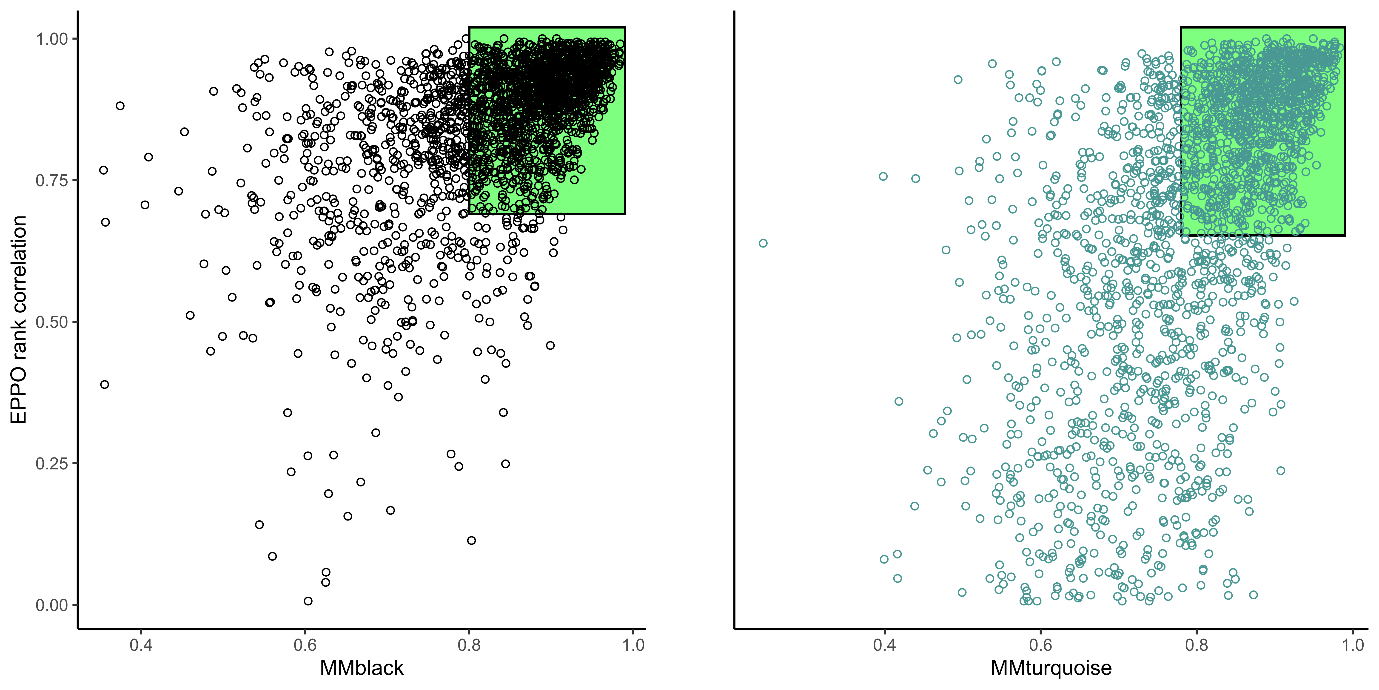

**Supplemental Figure 1.** Subsetting of the genes from WGCNA modules ‘black’ and ‘turquoise’ to select those most relevant to longevity acquisition. The genes are separated on the scatter plots’ x-axis based on module membership (MM), the correlation of a gene’s expression profile with the module eigengene of its respective WGCNA module, and on the y-axis by the correlation of their dry stage expression across genotypes with longevity ranks for those genotypes from the EPPO experiment. Only genes that fall in the area shaded green (top-right) were included in longevity motif enrichment analysis.

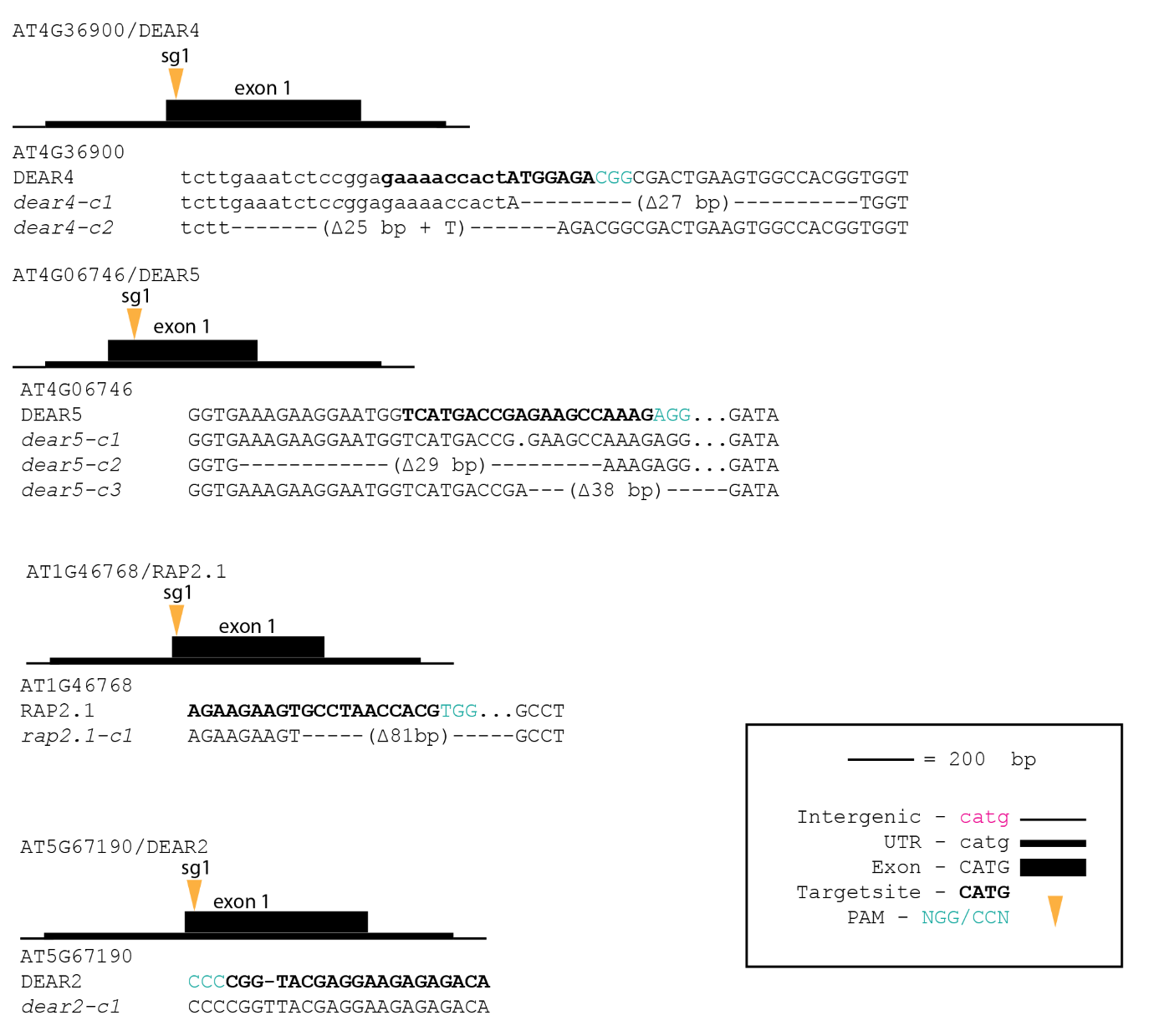

**Supplemental Figure 2.** Overview of CRISPR-Cas9 mutant construct generation. Sg1 - Single guide RNA 1.

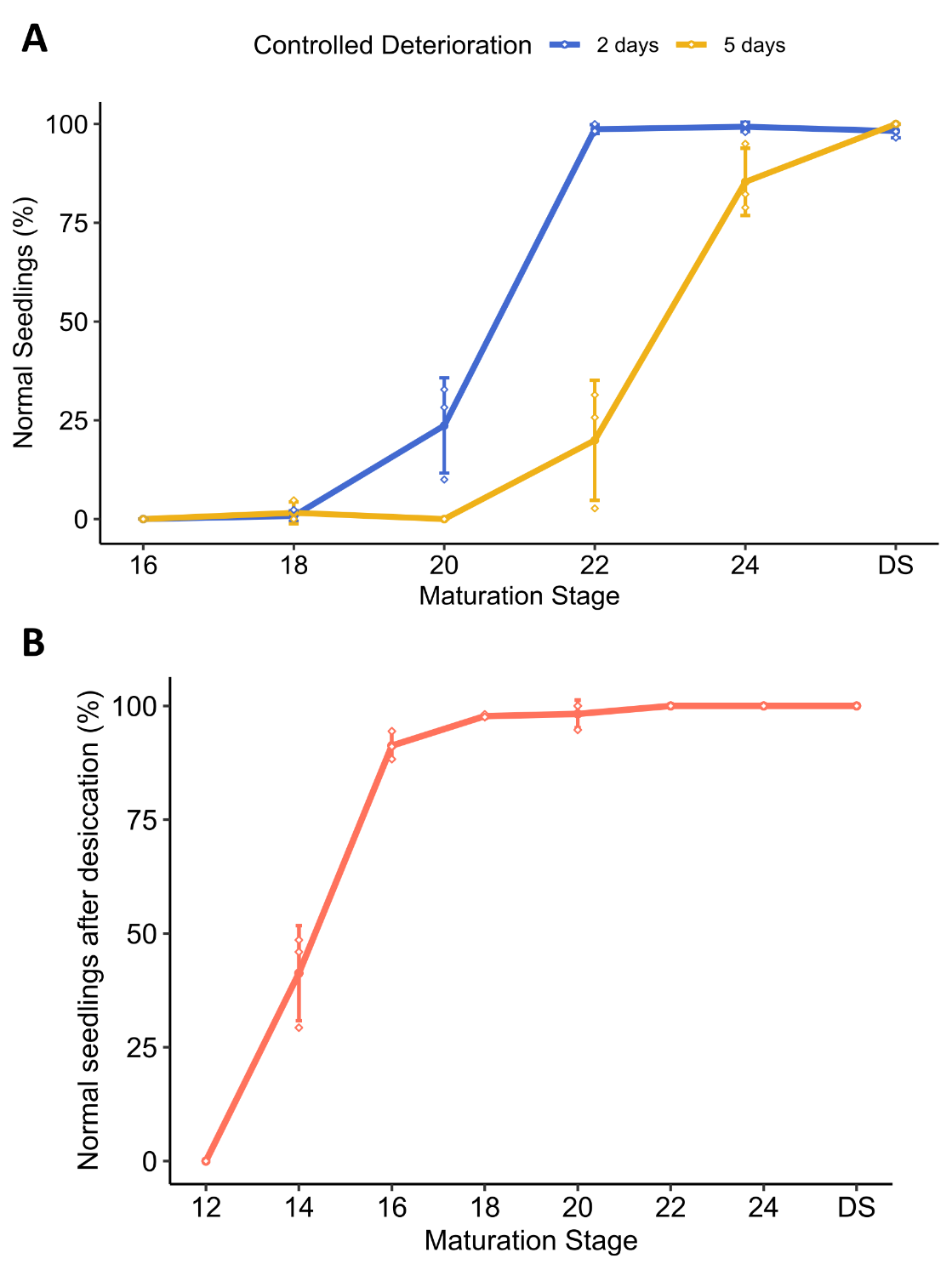

**Supplemental Figure 3.** Physiological characterization of longevity (A) and desiccation tolerance (DT) (B) acquisition during Col-0 seed maturation. A - Controlled deterioration test (CDT) was performed on maturing seeds every two days from 16 days after pollination (DAP) up to the dry seed (DS) stage. Seeds were stored for 2 days or 5 days at 80−85 % relative humidity (RH) and 40 °C prior to germination (n = 4). B – DT assessment was performed on maturing seeds every two days from 12 days after pollination (DAP) up to the dry seed (DS) stage. Seeds were desiccated for 48 hours at 22 °C and 30% relative humidity (RH) in the dark prior to germination (n = 4). Control consisted of seeds not desiccated and directly submitted to germination. DT percentage was calculated as the difference in the percentage of normal seedlings after 7 days between desiccated and non-desiccated samples. For both A-B, germination was performed on 10 μM GA_4+7_ and 10 mM KNO_3_ at 22 °C under continuous light.

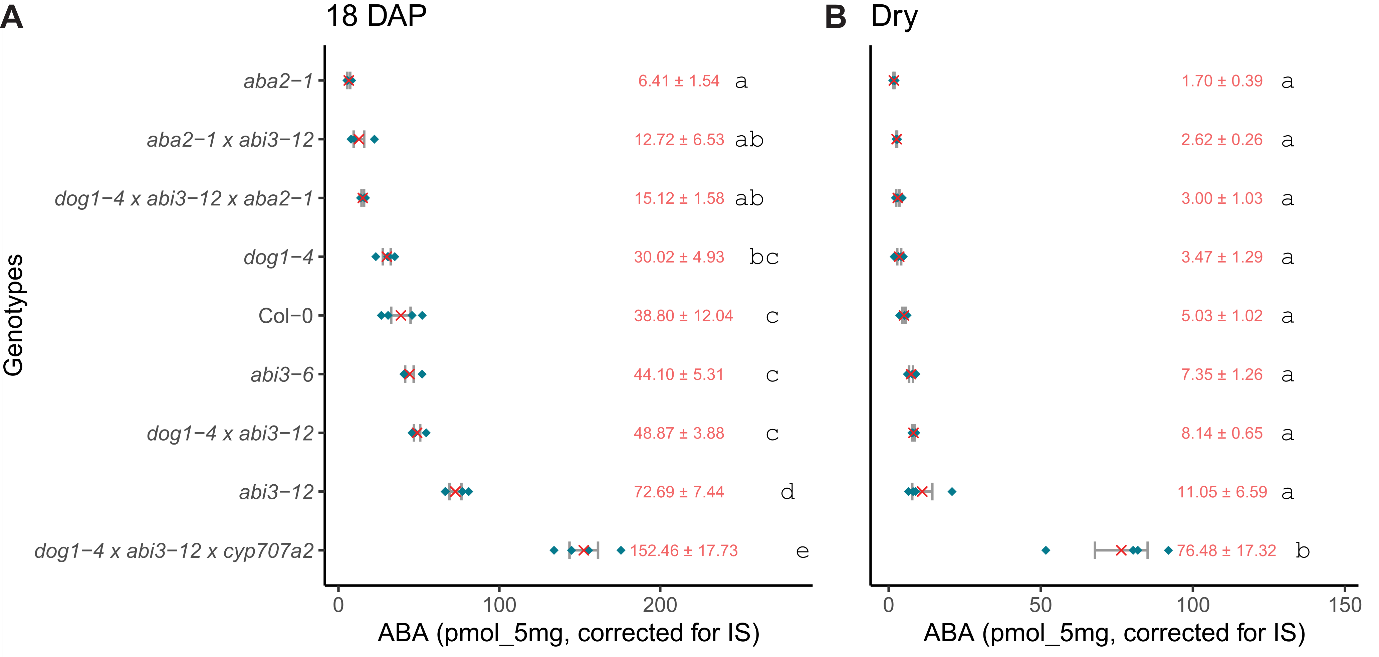

**Supplemental Figure 4.** ABA content in (A) 18 DAP and (B) dry seeds. Values are normalised by 5 mg of seeds and internal standard (IS) by LC-MS/MS, (n = 4). ANOVA was conducted to test for differences in mean ABA content across genotypes, pairwise comparisons were made using Tukey’s HSD post-hoc test. In the figure, a compact letter display illustrates statistically similar means: genotypes sharing the same letter in panel A or B have means that are not significantly different. Specifically, a shared letter indicates a Tukey HSD p-value above the significance threshold of 0.05.

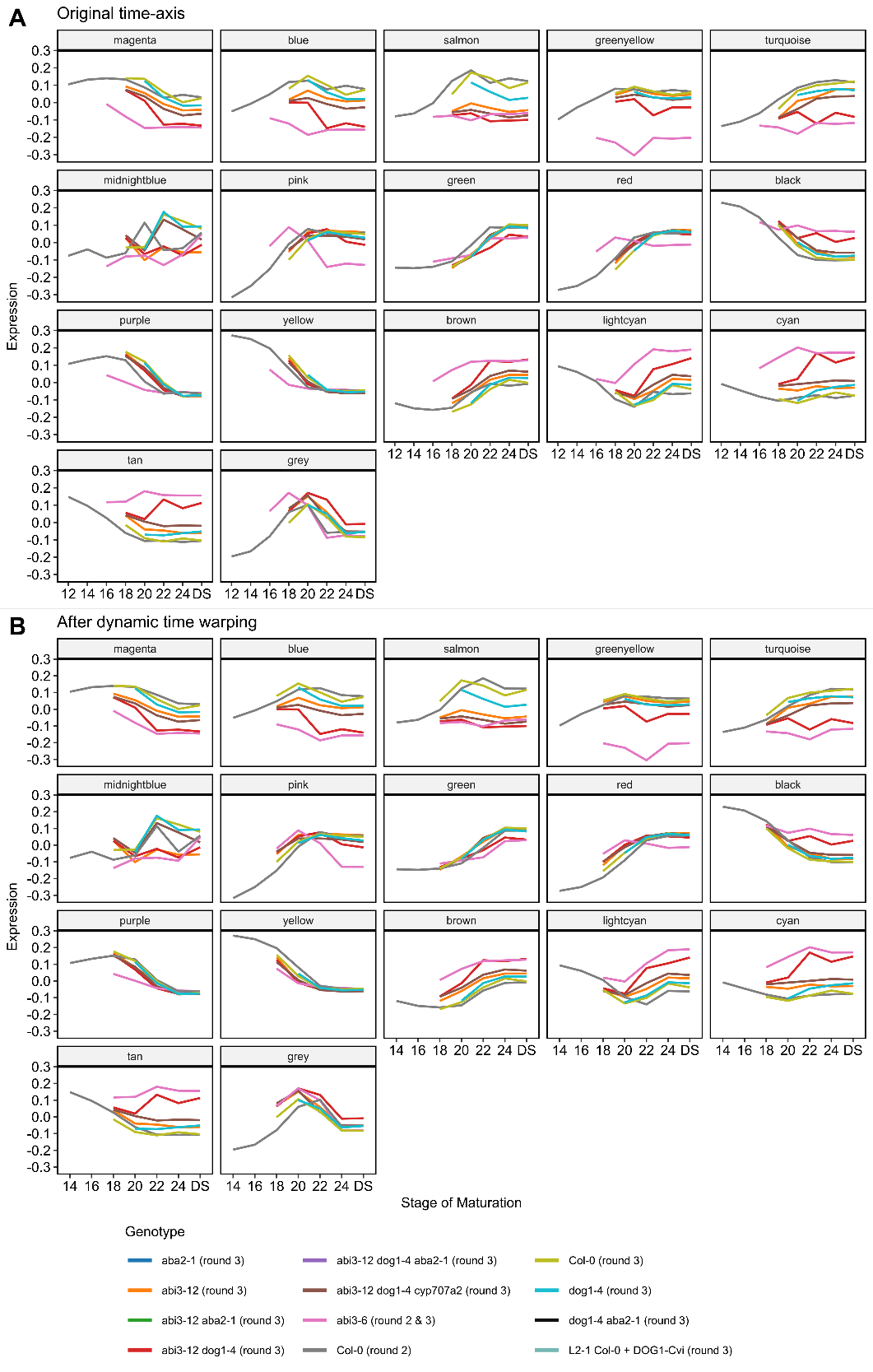

**Supplemental Figure 5.** Effect of dynamic time warping on module eigengenes (MEs). A. MEs without warping time axis. B. MEs with warped time axis. Time warping was used to adjust time point labels of round 2 to round 3, only Col-0 and *abi3-6* samples are affected. Adjustment of time axis labels for round 2 samples was as follows: +2 days for each sample from 12 to 22 days. This figure shows a better agreement of the ME profiles of Col-0 round 2 (grey) and Col-0 round 3 (lightgreen), clearly noticeable when comparing for the black module, before and after time-warping. Improved alignment of time-series for abi3-6 round 2 samples is noticeable when comparing the ME of *abi3-6* (pink) with the ME of *abi3-12 dog1-4 (red)*, for example for the turquoise module, before and after time-warping.

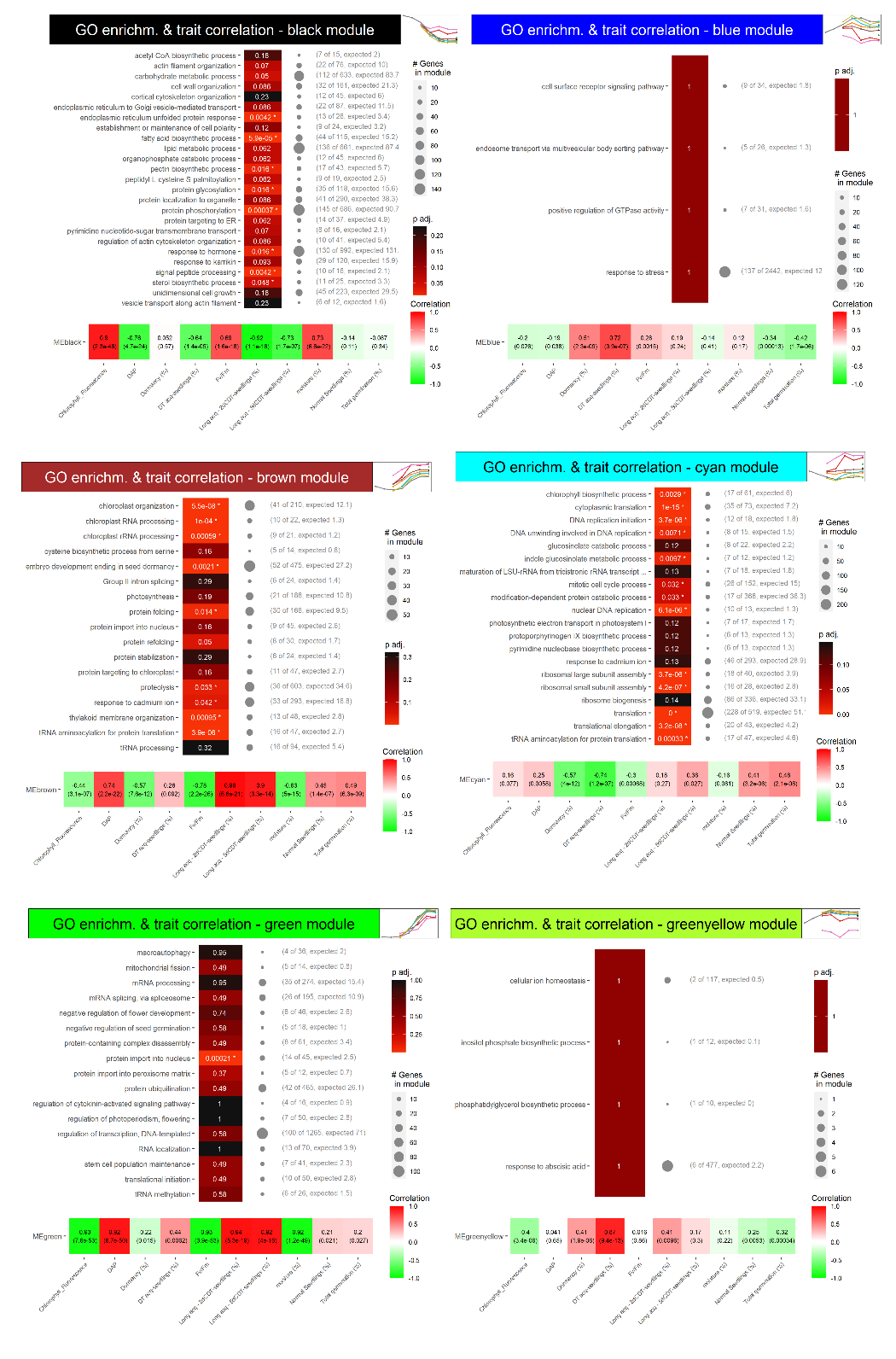

**Supplemental Figure 6 – Part I**. Overview of significantly enriched GO terms and trait correlations for modules obtained from WGCNA analysis. The selection of GO terms includes all biological processes with a significant enrichment (padj <0.05). A column with grey text displays the number of genes in each module annotated with a GO term, as well as the total number of genes annotated with the term in the background population. The expected number of genes associated with each GO term is shown based on the total number of genes in the module and the prevalence of the term in the background population. At the bottom of each graph, a heatmap is presented depicting the Pearson correlation between the eigengene of each module and a set of traits. The p−value corresponding to the correlation shown in parentheses.

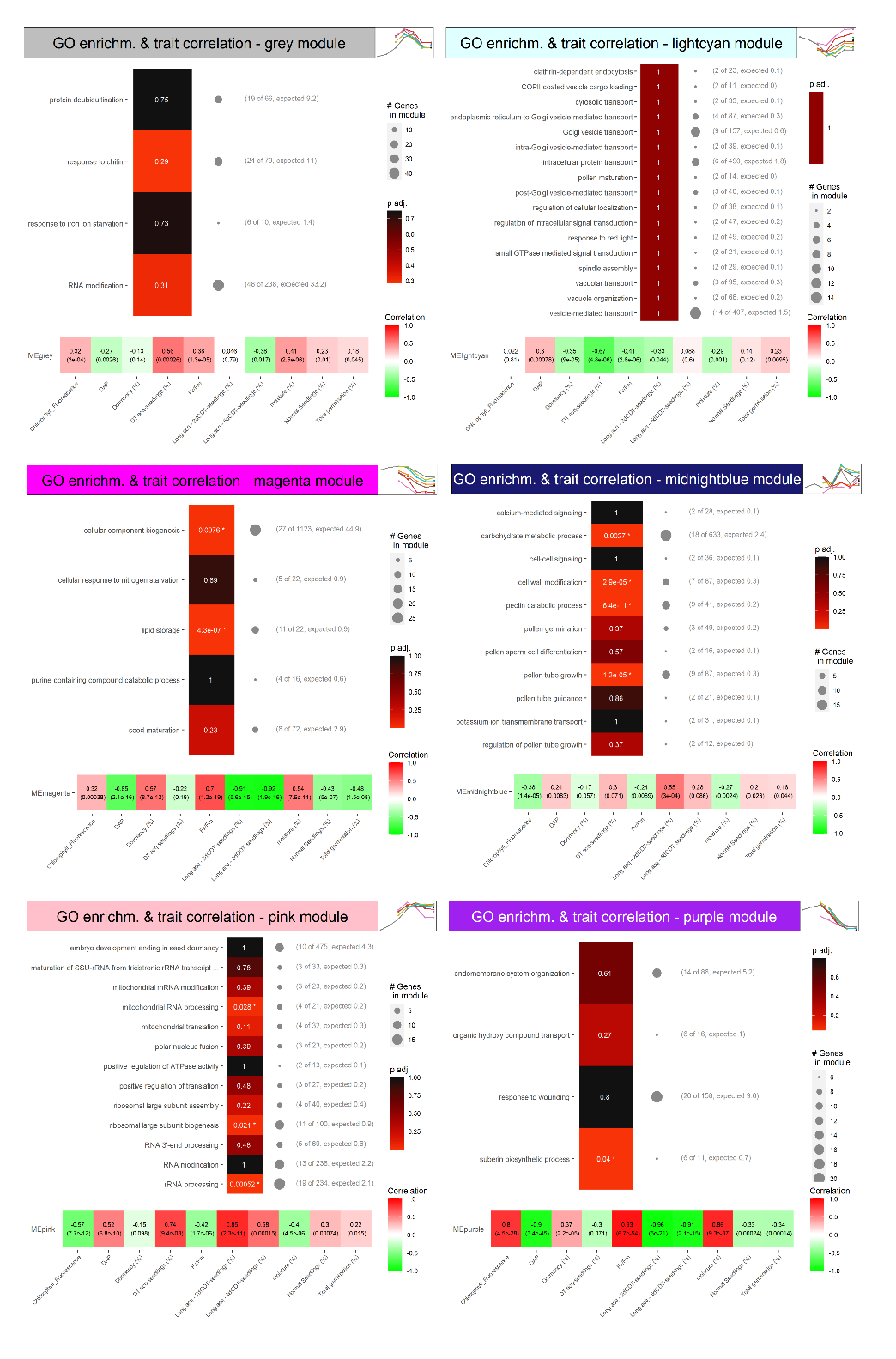

**Supplemental Figure 6 – Part II**. Overview of significantly enriched GO terms and trait correlations for modules obtained from WGCNA analysis. The selection of GO terms includes all biological processes with a significant enrichment (padj <0.05). A column with grey text displays the number of genes in each module annotated with a GO term, as well as the total number of genes annotated with the term in the background population. The expected number of genes associated with each GO term is shown based on the total number of genes in the module and the prevalence of the term in the background population. At the bottom of each graph, a heatmap is presented depicting the Pearson correlation between the eigengene of each module and a set of traits. The p−value corresponding to the correlation shown in parentheses.

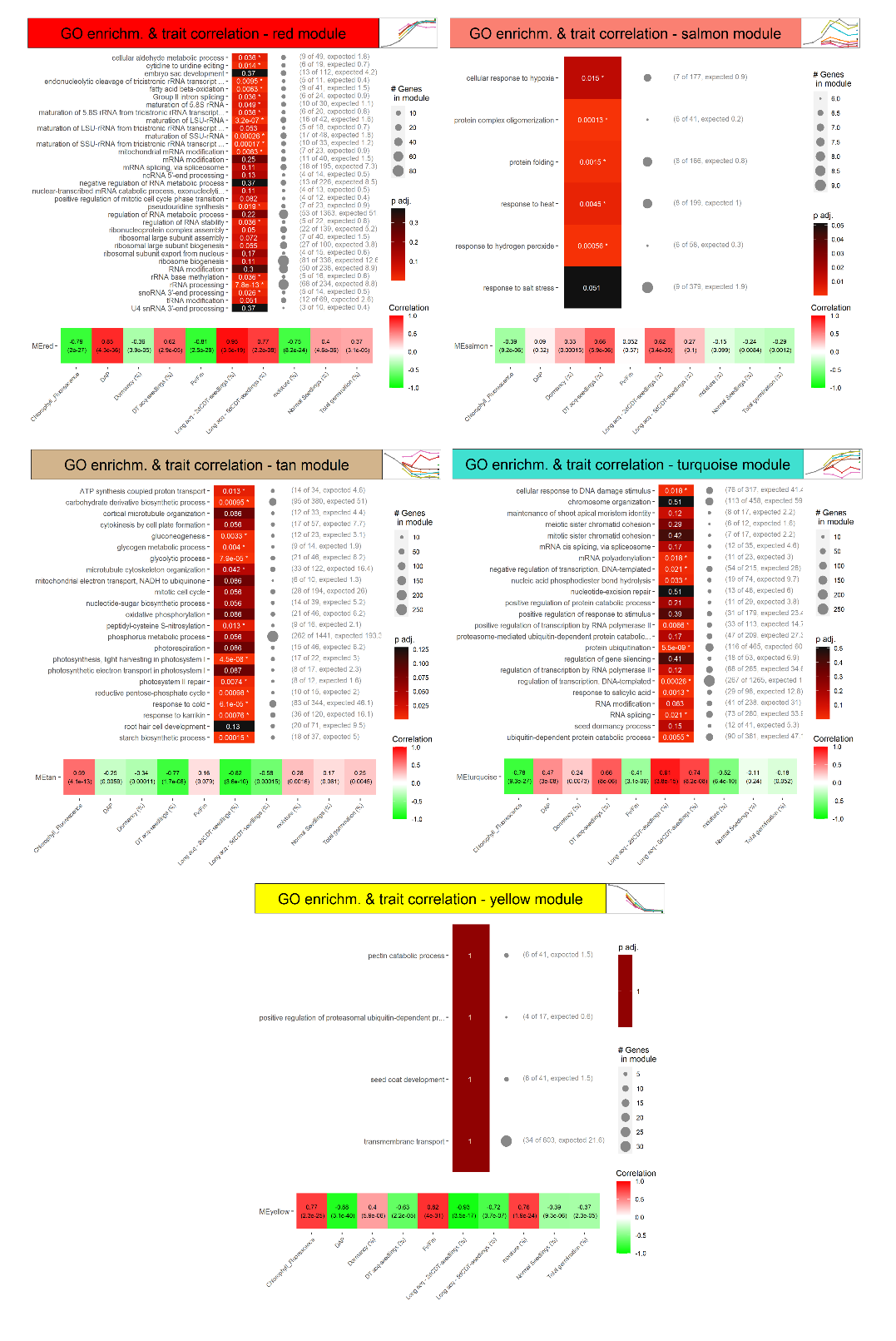

**Supplemental Figure 6 – Part III**. Overview of significantly enriched GO terms and trait correlations for modules obtained from WGCNA analysis. The selection of GO terms includes all biological processes with a significant enrichment (padj <0.05). A column with grey text displays the number of genes in each module annotated with a GO term, as well as the total number of genes annotated with the term in the background population. The expected number of genes associated with each GO term is shown based on the total number of genes in the module and the prevalence of the term in the background population. At the bottom of each graph, a heatmap is presented depicting the Pearson correlation between the eigengene of each module and a set of traits. The p−value corresponding to the correlation shown in parentheses.

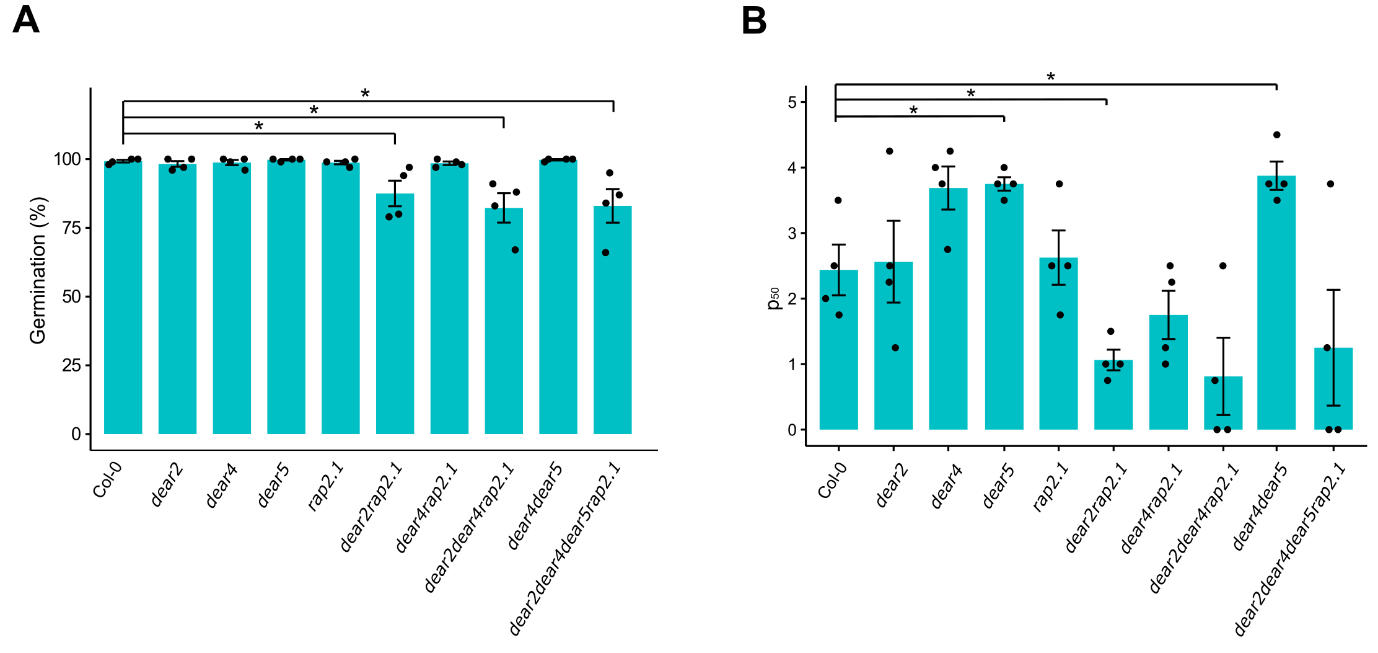

**Supplemental Figure 7**. A - Percentage of germination of mature dry seeds stored under natural conditions (50%RH, 20°C) for 47 days from Col-0 and multiple CRISPR-Cas9 mutants germinated in the presence of 10 μM GA_4+7_ and 10 mM KNO_3_. B - Half viability time (number of days to lose 50% of seed viability) of seeds submitted to CDT. Asterisks in A and B indicate significant differences (p<0.05) based on a Wilcoxon Sum Rank test and error bars indicate standard error (n = 4). The different CRISPR alleles and backgrounds are described in Supplemental Figure 3.
